# Tax1bp1 enhances bacterial virulence and promotes inflammatory responses during *Mycobacterium tuberculosis* infection of alveolar macrophages

**DOI:** 10.1101/2024.12.16.628616

**Authors:** Jeffrey Chin, Nalin Abeydeera, Teresa Repasy, Rafael Rivera-Lugo, Gabriel Mitchell, Vinh Q Nguyen, Weihao Zheng, Alicia Richards, Danielle L Swaney, Nevan J Krogan, Joel D Ernst, Jeffery S Cox, Jonathan M Budzik

## Abstract

Crosstalk between autophagy, host cell death, and inflammatory host responses to bacterial pathogens enables effective innate immune responses that limit bacterial growth while minimizing coincidental host damage. *Mycobacterium tuberculosis* (*Mtb*) thwarts innate immune defense mechanisms in alveolar macrophages (AMs) during the initial stages of infection and in recruited bone marrow-derived cells during later stages of infection. However, how protective inflammatory responses are achieved during *Mtb* infection and the variation of the response in different macrophage subtypes remain obscure. Here, we show that the autophagy receptor Tax1bp1 plays a critical role in enhancing inflammatory cytokine production and increasing the susceptibility of mice to *Mtb* infection. Surprisingly, although Tax1bp1 restricts *Mtb* growth during infection of bone marrow-derived macrophages (BMDMs) (Budzik *et al.* 2020) and terminates cytokine production in response to cytokine stimulation or viral infection, Tax1bp1 instead promotes *Mtb* growth in AMs, neutrophils, and a subset of recruited monocyte-derived cells from the bone marrow. Tax1bp1 also leads to increases in bacterial growth and inflammatory responses during infection of mice with *Listeria monocytogenes*, an intracellular pathogen that is not effectively targeted to canonical autophagy. In *Mtb-*infected AMs but not BMDMs, Tax1bp1 enhances necrotic-like cell death early after infection, reprogramming the mode of host cell death to favor *Mtb* replication in AMs. Tax1bp1’s impact on host cell death is a mechanism that explains Tax1bp1’s cell type-specific role in the control of *Mtb* growth. Similar to *Tax1bp1-*deficiency in AMs, the expression of phosphosite-deficient Tax1bp1 restricts *Mtb* growth. Together, these results show that Tax1bp1 plays a crucial role in linking the regulation of autophagy, cell death, and pro-inflammatory host responses and enhancing susceptibility to bacterial infection.

**Author Summary:** Although macrophages are the first innate immune cells to encounter *Mycobacterium tuberculosis* during infection, *M. tuberculosis* has evolved the ability to persist in them. Recent studies highlight that some types of macrophages are more permissive to *M. tuberculosis* replication and survival than others, but the mechanisms for cell type-specific differences in *M. tuberculosis* growth are only beginning to be understood. We found that the host factor, Tax1bp1 (Tax-1 binding protein 1), supports *M. tuberculosis* growth during animal infection and in specific subsets of innate immune cells, including alveolar macrophages while restricting *M. tuberculosis* in bone marrow-derived macrophages. We also found that Tax1bp1 has a similar phenotype in enhancing the pathogenesis of another intracellular pathogen, *Listeria monocytogenes.* Compared to bone marrow-derived macrophages, in alveolar macrophages, Tax1bp1 enhances the release of inflammatory mediators and leads to necrotic-like host cell death, which is known to enhance *M. tuberculosis* growth. Phosphorylation of Tax1bp1 in alveolar macrophages promotes *M. tuberculosis* growth. Our research highlights that Tax1bp1 is a host target for host-directed therapy against *M. tuberculosis* and controls host responses to *M. tuberculosis* in a cell type-specific manner.

## Introduction

*Mycobacterium tuberculosis* (*Mtb*), the causative agent of tuberculosis, has evolved the ability to circumvent host innate antimicrobial responses and survive within our immune cells (1,2). Understanding the host factors that limit antimicrobial immune responses to *Mtb* is critical for developing new host-directed antimicrobial therapies (3,4). The persistently high number of deaths from tuberculosis (1.13 million in 2023 (5)) highlights the need for new anti-TB treatments.

After inhalation of *Mtb*, alveolar macrophages (AMs) are the first immune cells to become infected with *Mtb* and provide a replicative niche for intracellular bacteria (6–8). To disseminate to other organs, *Mtb* must spread from AMs to different cell types, such as recruited monocyte-derived macrophages (6). In the mouse model of infection, this dissemination occurs at approximately 14-25 days post-infection upon recruitment of monocytes and their differentiation into macrophages in the lung parenchyma (9).

Ultimately, live bacteria are transported from the lungs via lymphatics to the draining lymph nodes (9). In *Mtb-*infected lungs, monocytes have been shown to differentiate into two subsets that differ in their ability to control *Mtb* (10). The first subset is the CD11c^lo^ subset (MNC1, mononuclear cell subset 1), also known as recruited macrophages (11–13). The second subset is the CD11c^hi^ subset (MNC2), formerly known as dendritic cells but considered more closely related to macrophages (9,12–14). AMs, which are embryonically derived from the yolk sac, and monocyte-derived macrophages originating from circulating monocytes of bone marrow origin (15) exhibit divergent transcriptional responses against *Mtb* (8). AMs are impaired at mounting antibacterial responses against *Mtb* which lead to significant *Mtb* growth differences when compared to murine bone marrow-derived macrophages (BMDMs) (8). However, the host factors mediating different rates of *Mtb* growth in AMs compared to BMDMs are poorly understood.

In various macrophage subtypes, immune responses to *Mtb* are generated by the detection of *Mtb* lipoproteins and lipoglycans by macrophage surface receptors (*e.g.* TLR2 (16,17), Dectin 1 (18)), which mediate ERK (extracellular-signal regulated kinase) and NF-κB inflammatory signaling (19). Cytosolic sensors also recognize *Mtb* nucleic acids released during phagosomal perforation (20) and engage several downstream signaling pathways, each of which can have opposing impacts on *Mtb* growth. For example, cytosolic sensing of *Mtb* DNA by cGAS (cyclic GMP–AMP synthase) triggers type I interferon (IFN) production and autophagy (21). Autophagy is a protective response against *Mtb* that targets it for ubiquitylation and degradation in the lysosome (22–25). Conversely, type I IFN is a cytokine that protects against viruses but can be co-opted by *Mtb* to promote its growth (26–28). Another important cytosolic sensor for *Mtb* is the RIG (Retinoic acid-inducible gene I)-I-like pathway. Like the cGAS pathway, the RIG-I-like pathway’s cytosolic sensing of *Mtb* RNA induces host responses with opposing effects on *Mtb* growth. RIG-I induces apoptosis, a mode of cell death that leads to *Mtb* growth restriction, while also triggering pro-bacterial type I IFN and limiting NF-κB cytokine production such as TNF-α, IL-1β, and IL-6. The impacts of NF-κB-regulated cytokines can be pro-or anti-bacterial (29,30). For instance, TNF-α can restrict *Mtb* by activating phagocytes but, in excess, can enhance *Mtb* growth by mediating tissue damage (31–34). These *Mtb*-mediated signaling pathways are regulated by post-translational modifications through cascade signaling protein phosphorylation (20,22,25,35). Thus, *Mtb* infection triggers post-translationally regulated host responses that can have pro-or anti-bacterial effects. Nevertheless, we lack a complete understanding of host factors that drive these responses to thwart *Mtb* growth.

Tax1bp1 is an autophagy receptor at the nexus of multiple key immune responses critical for pathogen control. Tax1bp1 regulates inflammation and blocks apoptosis in response to cytokine stimulation, vesicular stomatitis virus (VSV), and Sendai virus infections by terminating NF-κB and RIG-Isignaling (36–42). In infected Tax1bp1-deficient mice, respiratory syncytial virus (RSV) replication is decreased, whereas cytokine responses are enhanced (37). The anti-inflammatory function of Tax1bp1 has also been shown to impact non-infectious diseases through abrogating development of chemically-induced hepatocellular cancer (38), age-dependent dermatitis, and cardiac valvulitis (36). Tax1bp1 promotes selective autophagy that mediates lysosomal degradation of pathogens, such as *Mtb* in BMDMs (43) and *Salmonella enterica* Typhimurium (44). Additionally, Tax1bp1 mediates the clearance of aggregated neuronal proteins involved in neurodegenerative disease (45). Thus, Tax1bp1 has an anti-inflammatory function in several contexts, but the impact of Tax1bp1 *in vivo* during intracellular bacterial infection has hitherto not been described.

Our previous work showed that the autophagy receptor, Tax1bp1, is phosphorylated during *Mtb* infection of BMDMs (43). Tax1bp1 restricts *Mtb* growth during *ex vivo* infection of BMDMs presumably because of the role of Tax1bp1 in antibacterial autophagy (43). Notably, Tax1bp1 does not significantly change the levels of NF-κB-regulated cytokines during *Mtb* infection of BMDMs (43). To expand our understanding of Tax1bp1’s function in other critical innate immune cell types and the presence of the complete immune system, here we employed the mouse infection models for *Mtb* and the intracellular pathogen *Listeria monocytogenes*. Surprisingly, this led to the discovery that Tax1bp1 promotes *Mtb* and *Listeria* growth during animal infection and enhances *Mtb* growth in several innate immune cell types including AMs, in contrast to the role of Tax1bp1 in the restriction of *Mtb* growth in BMDMs. Furthermore, Tax1bp1 had a pro-inflammatory function during *Mtb* and *Listeria* animal infection, compared to its anti-inflammatory role in viral infections. We found that Tax1bp1 enhances necrotic-like cell death and inflammatory mediator release during AM but not BMDM infection, which is a mechanism by which Tax1bp1 leads to cell type-specific changes in *Mtb* growth. To our knowledge, we are the first to study the function of Tax1bp1 in the context of mouse infections with these pathogens. Our findings from the animal infection model led us to uncover new relevant phenotypes compared to those revealed by the BMDM infection model and revealed Tax1bp1’s negative impact on immunity to intracellular pathogens.

## Results

### Tax1bp1 promotes *Mtb* virulence and inflammatory cytokine responses *in vivo*

To assess the role of *Tax1bp1*’s contribution to controlling tuberculosis infection *in vivo*, we infected wild-type and Tax1bp1-deficient mice with virulent *M. tuberculosis* via the aerosol route and monitored the infection at different time points. First, we infected male and female wild-type and *Tax1bp1^−/−^* mice at a low dose and enumerated CFU at day one post-infection. This revealed no statistically significant difference in bacterial uptake (Figure 1A). Next, two independent mouse infections were performed on separate days to test the contribution of Tax1bp1 to *Mtb* growth (Figure 1B, Figure 1-figure supplement 1). On days 9 or 11, 21, and 50 post-infection, we harvested the lung, spleen, and liver for enumeration of bacterial CFU. In contrast to our previous results showing Tax1bp1 restricted *Mtb* growth in BMDMs infected *ex vivo* (43), in the mouse infection model, Tax1bp1 enhanced *Mtb* growth (Figure 1B, Figure 1-figure supplement 1). This unexpected difference manifests even after 11 days of infection, during the acute stage when *Mtb* replicates within AMs. *Mtb*’s improved growth was magnified in both peripheral organs, liver, and spleen, consistent with the differences in lung CFUs. These results contrast with the autophagy receptor function of Tax1bp1 since Tax1bp1 contributes to targeting *Mtb* to selective autophagy (43) and autophagy is required for controlling *Mtb* growth *in vivo* and *ex vivo* (22,46).

**Figure 1.**
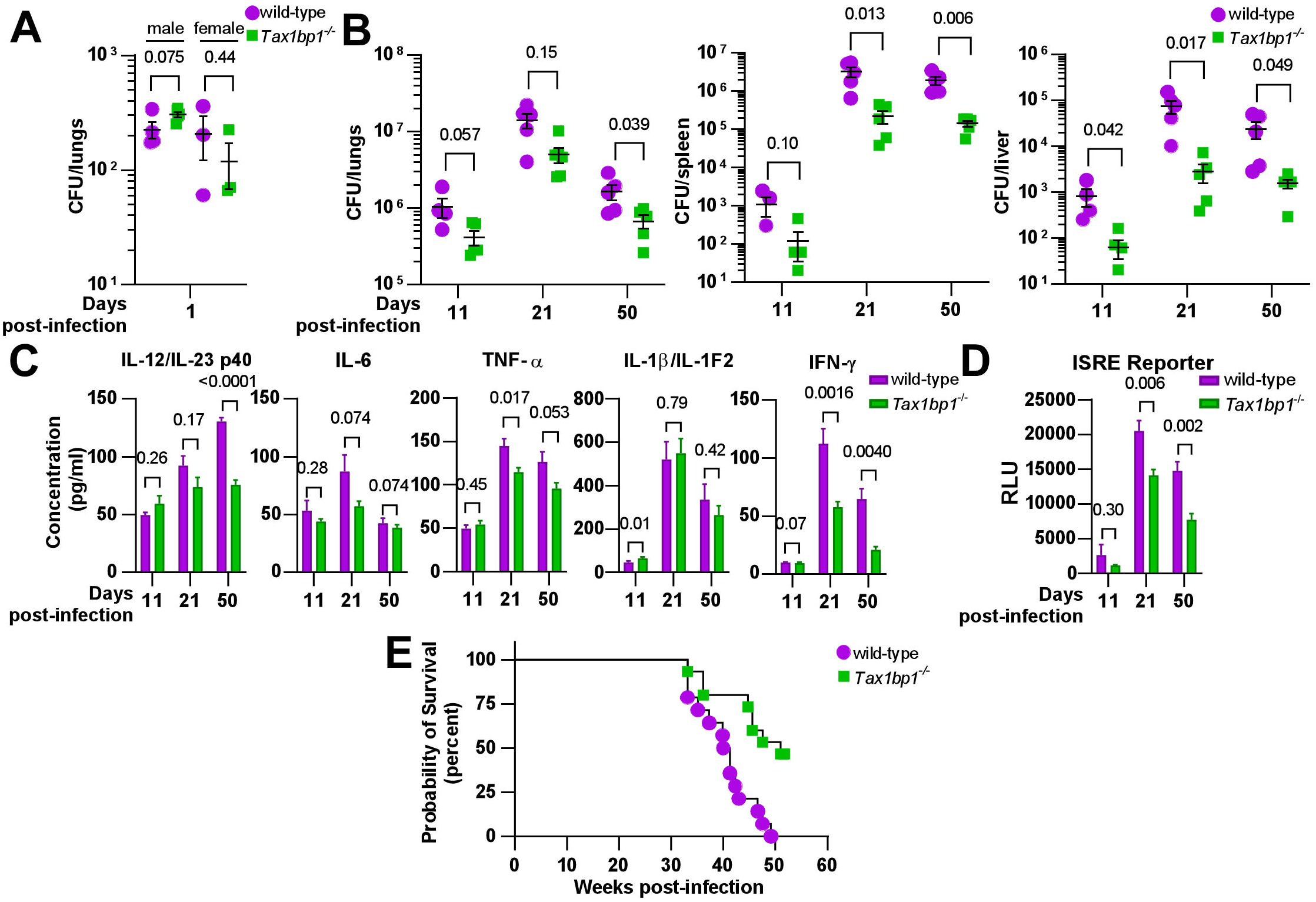
Tax1bp1 enhances *M. tuberculosis* virulence and inflammatory cytokine responses during mouse aerosol infection. (A) Male and female mice were infected by the aerosol route with a mean *Mtb* CFU of 240 as determined by CFU enumeration from lung homogenates at 1-day post-infection. (B) Additional mice were euthanized at 11-, 21-, and 50 days post-infection for CFU enumeration. Results are the mean ± SEM from lung homogenates of 5 infected mice. (C) Cytokine levels from infected lung homogenates at 11-, 21-, and 50-days post-infection were measured by ELISA. Results are the mean ± SEM from five samples. (D) Levels of type I and II interferon-induced JAK/STAT signaling were measured by luminescence in relative light units (RLUs) from infected lung homogenates by the ISRE assay. Results are the mean ± SEM from five samples. Brackets indicate p-values from t*-*test comparisons. (E) Infected mice were monitored for death or 15% loss of maximum body weight, at which point they were euthanized. Log-rank (Mantel-Cox) test and Gehan-Breslow-Wilcoxon comparison test p-values for survival were 0.0008 and 0.0047, respectively.

Because Tax1bp1 has a known role in mediating inflammatory responses, we hypothesized that Tax1bp1 might be regulating inflammatory responses during *Mtb* infection. Indeed, analysis of a panel of pro-inflammatory cytokines from the lungs of infected mice revealed that, consistent with this idea, Tax1bp1 increased levels of IL-6, TNF-α, IL1-β, and IL-12/IL-23 p40 (Figure 1C, Figure 1-figure supplement 1). Type I and II IFN are particularly important for controlling *Mtb* infection (21,28,47). Therefore, we measured interferon levels using a type I and II IFN reporter cell line (ISRE) and a type II IFN (IFN-γ) by ELISA (Figure 1C, D). Consistent with other pro-inflammatory cytokines, we found that wild-type mice had significantly higher levels of type I and II IFN in the lungs compared to *Tax1bp1*^−/−^ mice (Figure 1C, D). We also carried out survival studies and found that Tax1bp1 contributes to mortality during *Mtb* infection (Figure 1E). Although Tax1bp1 contributes to inflammatory cytokine synthesis during *Mtb* infection, microscopic examination of infected lung tissue did not reveal any significant differences in the cellular infiltrate of the lungs as reflected by lesion severity or tissue necrosis (Figure 1-figure supplement 2A, B) or by neutrophil recruitment reflected by myeloperoxidase staining (Figure 1-figure supplement 2C, D). During infection of BMDMs, we previously observed that Tax1bp1 reduces ubiquitin colocalization with *Mtb* (43). Thus, we sought to determine whether ubiquitin recruitment was regulated during infection *in vivo.* While Tax1bp1 led to a slight decrease in ubiquitin and *Mtb* colocalization in infected lung tissue samples at 50 days post-infection, this did not reach statistical significance (Figure 1-figure supplement 3). Our results suggest that Tax1bp1 amplifies host-detrimental inflammatory responses, which predominate over cell-intrinsic control of *Mtb* replication by autophagy in macrophages (48) and contribute to host susceptibility and mortality.

### Tax1bp1 enhances *Listeria monocytogenes* growth, microabscess formation, and host inflammatory cytokine synthesis

We then turned to a different intracellular bacterial pathogen, *Listeria monocytogenes*, which has potent autophagy-inhibiting mechanisms to avoid antibacterial autophagy (49–51). Using *Listeria* allowed us to assess the role of Tax1bp1 in inflammatory responses in isolation from its role in autophagy. As shown in Figures 2A and 2B, wild-type *Listeria* grew as well in both BMDMs and peritoneal macrophages harvested from wild-type and *Tax1bp1*^−/−^ mice, indicating there is no defect in the ability of *Listeria* to replicate in *Tax1bp1*^−/−^ macrophages *ex vivo*. In contrast, when we infected mice with *Listeria* via the intravenous (IV) route, we saw that over a 48-hour time course, *Tax1bp1^−/−^* mice were remarkably resistant to *Listeria* growth (Figure 2C). Consistent with Tax1bp1 playing a role early in the *Listeria* infection, we observed a statistically significant difference in *Listeria* CFU in the spleen, but not in the liver, at 4 hours post-infection (Figure 2D). This is consistent with the role of Tax1bp1 being manifested at the earliest stages of infection. To confirm the reproducibility of this growth phenotype, we repeated the *Listeria* mouse infections and observed similar results (Figure 2-figure supplement 1).

**Figure 2.**
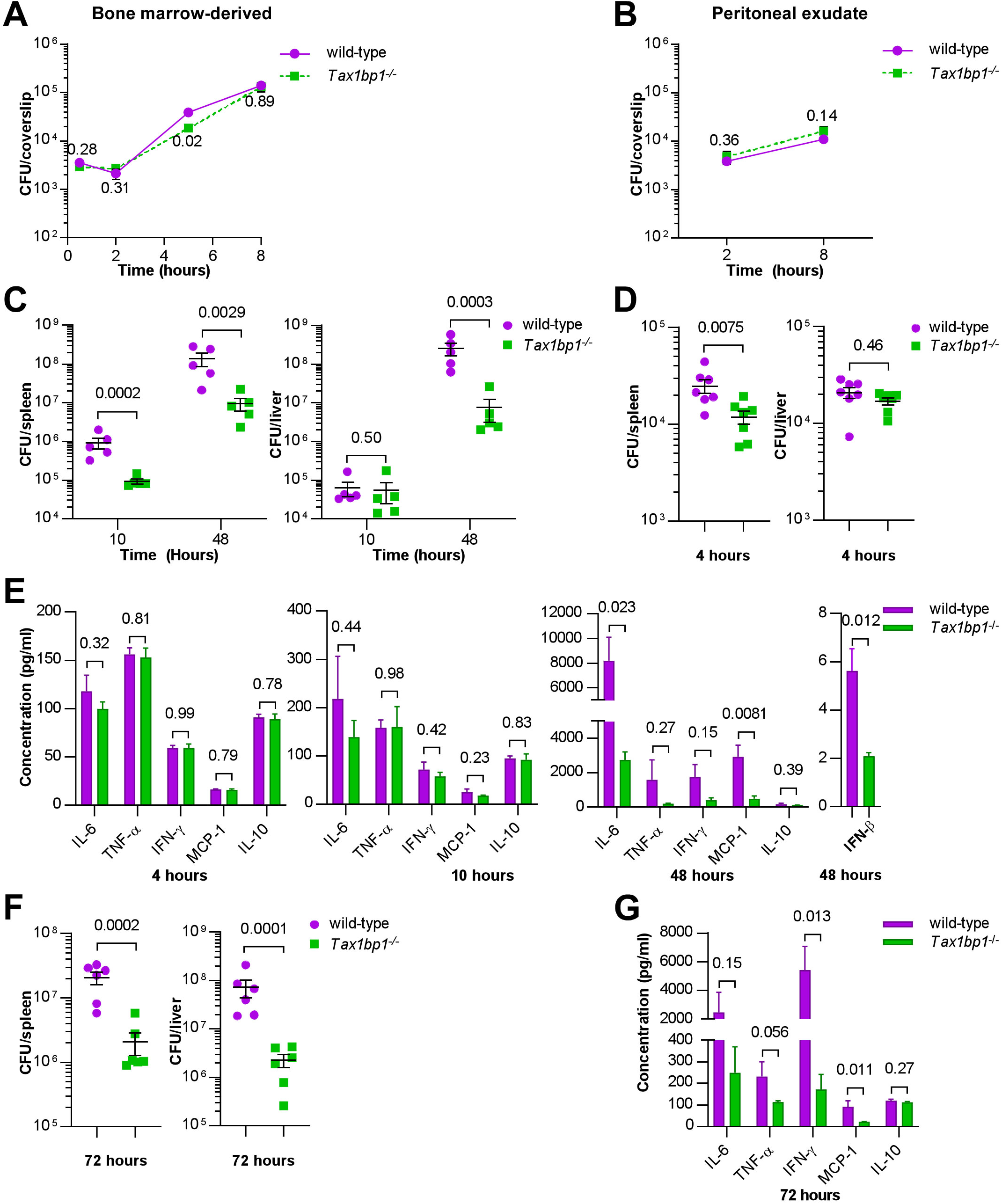
Tax1bp1 contributes to *Listeria monocytogenes* virulence and growth during murine but not *ex vivo* cellular infections. (A and B) BMDMs or peritoneal exudate cells were infected with *L. monocytogenes,* and CFU were counted at 30 minutes, 2-, 5-, or 8 hours post-infection. Results are the mean ± SEM from three technical replicate samples. The p-values from t-test comparisons are shown. (C and D) CFU from spleen and liver homogenates from mice intravenously infected with *L. monocytogenes* were enumerated at 4-, 10-, or 48-hours post-infection. Results are the mean ± SEM from five mice. Brackets indicate p-values from t-test comparisons. (E & G) Cytokine levels were measured from the serum of mice infected with *L. monocytogenes* at 4-, 10-, and 48-hours post-infection by cytometric bead array (IL-6, TNF-α, IFN-γ, MCP-1, IL-10) or ELISA (IFN-β). Results are mean ± SEM from five samples. Brackets indicate p-values from t*-*test comparisons. (F) Spleen and liver homogenates from mice intraperitoneally infected with *L. monocytogenes* were enumerated for CFU 72 hours post-infection. Results are the mean ± SEM from five mice. Brackets indicate p-values from t-test comparisons. CFU data were logarithmically transformed prior to statistical analysis.

We measured inflammatory cytokine levels in the serum of the IV-infected mice and found that during the first 4– and 10 hours, inflammatory cytokines were low and were not significantly different between the wild-type and *Tax1bp1^−/−^* mice (Figure 2E). At 48 hours, Tax1bp1 significantly increased IL-6, TNF-α, IFN-γ, IFN-β and MCP-1 in the serum, indicating Tax1bp1 is required for augmenting pro-inflammatory cytokines and type I interferon (IFN-β) (Figure 2E). Infecting mice by the intraperitoneal route gave rise to very similar results, indicating that the route of infection is irrelevant to the infection outcome (Figure 2F). Indeed, Tax1bp1 increased CFU by approximately 1.5 logs in the liver and by 1 log in the spleen (Figure 2F). As with the IV infection, Tax1bp1 led to a considerable non-statistically significant increase in IL-6 production and a substantial increase in IFN-γ in the serum (Figure 2G).

Likewise, Tax1bp1 augmented MCP-1 levels in the serum (Figure 2G). Consistent with the increased bacterial load mediated by Tax1bp1, histological examination of tissues from infected mice revealed that Tax1bp1 increased the number of microabscesses and led to lymphoid depletion (Figure 2-figure supplement 2). Tax1bp1 also reduced the occurrence and severity of hepatocyte coagulative necrosis (Figure 2-figure supplement 2). The lack of splenic microabscesses and lymphoid depletion of *Tax1bp1^−/−^*mice may correlate with the decrease in pro-inflammatory cytokine levels noted in the serum of *Tax1bp1^−/−^* mice compared to wild-type mice. These results suggest that Tax1bp1 enhances inflammatory cytokine signaling and bacterial growth during animal infection with *Listeria monocytogenes*, as observed during *Mtb* infection.

### Tax1bp1 promotes *Mtb* growth in AMs, neutrophils, and recruited mononuclear cells *in vivo*

Since Tax1bp1 restricted *Mtb* growth in BMDMs infected *ex vivo* (43) but the opposite phenotype was observed during *in vivo* mouse infections, we hypothesized that Tax1bp1 promotes *Mtb* growth in other cell types. Following the transfer of *Tax1bp1^−/−^*mice from UC Berkeley to UC San Francisco and their rederivation, we infected wild-type and *Tax1bp1^−/−^* age– and sex-matched mice with a low dose of ZsGreen-expressing *Mtb* by the aerosol route. At 7– and 14 days post-infection, *Mtb* CFU were enumerated from aliquots of organ homogenates. Consistent with our previous results at UC Berkeley, Tax1bp1 promoted *Mtb* growth in rederived mice (Figure 3A). To test the hypothesis that Tax1bp1 enhances *Mtb* growth in distinct innate immune cells, lung cell suspensions were pooled from five mice of each genotype to obtain enough cells for downstream CFU analysis from sorted cell populations. After staining the pooled innate immune cells, we performed flow cytometry to sort CD11c^lo^ (MNC1) and CD11c^hi^ (MNC2) mononuclear cells, neutrophils (PMNs), and AMs using innate immune cell antibodies (Figure 3-figure supplement 1) (10). To account for differences in the number of cells sorted between genotypes, data were normalized by dividing the ZsGreen+ counts or CFU by the total number of each cell type sorted. Tax1bp1 increased the normalized ZsGreen+ counts in AMs, neutrophils, MNC1, and MNC2 (Figure 3B). We plated the sorted innate immune cells on agar plates in quadruplicate to determine the number of viable *Mtb* CFU in the sorted cells. Consistent with the ZsGreen+ counts, Tax1bp1 promoted *Mtb* growth by CFU counts in AMs by 11-fold and in PMNs by 6-fold at seven days post-infection (Figure 3C). At 14 days post-infection, the phenotype was more pronounced in AMs (43-fold increase) and less pronounced in PMNs (2-fold increase; Figure 3C). CFU were not detected from MNC1 and MNC2 at 7-days post-infection, consistent with a previous report (52). Although Tax1bp1 increases ZsGreen+ counts in MNC1 at 14-days post-infection, Tax1bp1 did not enhance *Mtb* CFU in MNC1 but did slightly in MNC2 (1.6-fold; Figure 3C). We performed a second independent aerosol infection to test the reproducibility of these findings and measure ZsGreen+ *Mtb* counts at an additional point 21 days post-infection. As observed in the first experiment, Tax1bp1 increased *Mtb* counts at the early time points in AMs, MNC2, and PMNs; however, at 21 days post-infection Tax1bp1 instead reduced *Mtb* ZsGreen+ counts in MNC1 (Figure 3-figure supplement 2). This suggests that Tax1bp1 can restrict *Mtb* growth in MNC1, consistent with our previous findings in BMDMs *ex vivo*. In summary, Tax1bp1 has a cell type-specific impact on *Mtb* growth with an overall effect of increasing *Mtb* growth in the major organs. In two independent aerosol infection experiments, Tax1bp1 enhances *Mtb* growth in AMs, PMNs, and MNC2, boosting *Mtb* growth in immune cells of various cell origins *in vivo*.

**Figure 3.**
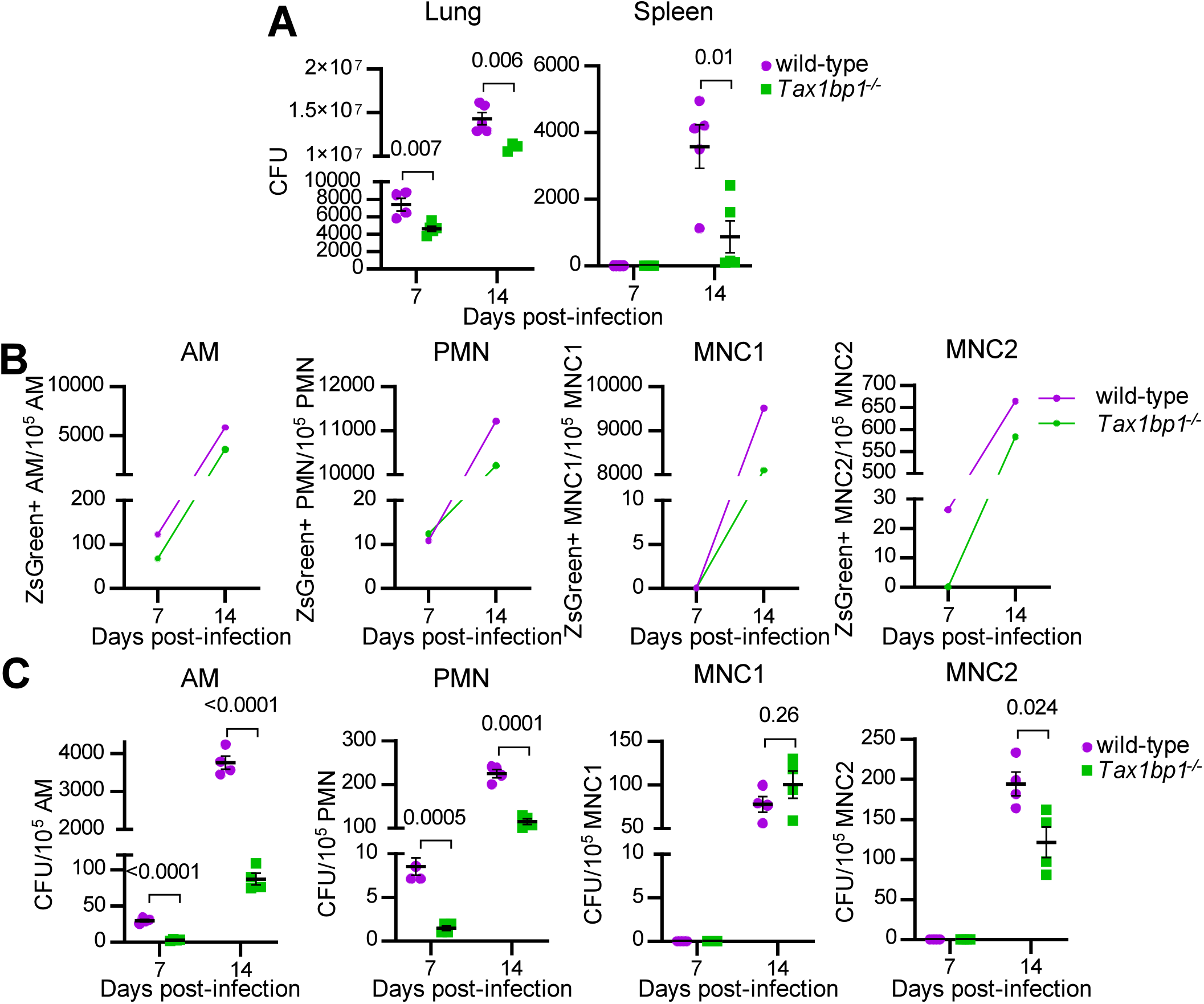
Tax1bp1 promotes *Mtb* growth in AMs, PMNs, and MNC2 following low-dose aerosol infection. Mice were infected with aerosolized *Mtb* expressing ZsGreen (calculated dose of 100 CFU per mouse), and five wild-type and 5 *Tax1bp1^−/−^*mice were euthanized at 7– and 14-days post-infection. (A) Lung and spleen homogenates from 5 wild-type and 5 *Tax1bp1^−/−^* mice at each time point were plated for CFU. (B) Lung cells were pooled and stained for AMs, neutrophils (PMNs), and recruited monocyte 1 and 2 subsets (MNC1, 2). ZsGreen-positive innate immune cell subsets were quantified by analytical flow cytometry. (C) Innate immune cells were sorted. The sorted cells were plated for *Mtb* CFU in quadruplicate. Data were normalized to the number of cells sorted. SEM and p-values from the t-test are displayed.

To test whether Tax1bp1 also enhances *Mtb* growth in AMs infected *ex vivo*, we obtained murine AMs by bronchoalveolar lavage (BAL), discarded the suspension cells, and cultured the remaining adherent AMs. We infected the AMs with either luminescent or wild-type *Mtb* in the presence or absence of IFN-γ, a cytokine that activates macrophage antibacterial host responses (53). As reflected by changes in luminescence (Figure 4A), Tax1bp1 began to promote *Mtb* growth at two days post-infection in the presence or absence of IFN-γ. Similarly, Tax1bp1 augmented *Mtb* growth as measured by CFU (Figure 4B). This data implicates Tax1bp1 in enhancing *Mtb* growth during AM infection and is consistent with our *in vivo* studies, although it is in sharp contrast to our results in *Mtb-*infected BMDMs (43) and MNC1. Therefore, we focused on understanding Tax1bp1’s function in AMs as this model better represents the overall impact of Tax1bp1 on real-world infections and in most innate immune cell types. Even though Tax1bp1’s phenotype in BMDMs does not model the phenotype in several other cell types (*i.e.,* AMs, MNC2, and PMNs), we took advantage of Tax1bp1’s phenotype in BMDMs by using BMDMs as a control.

**Figure 4.**
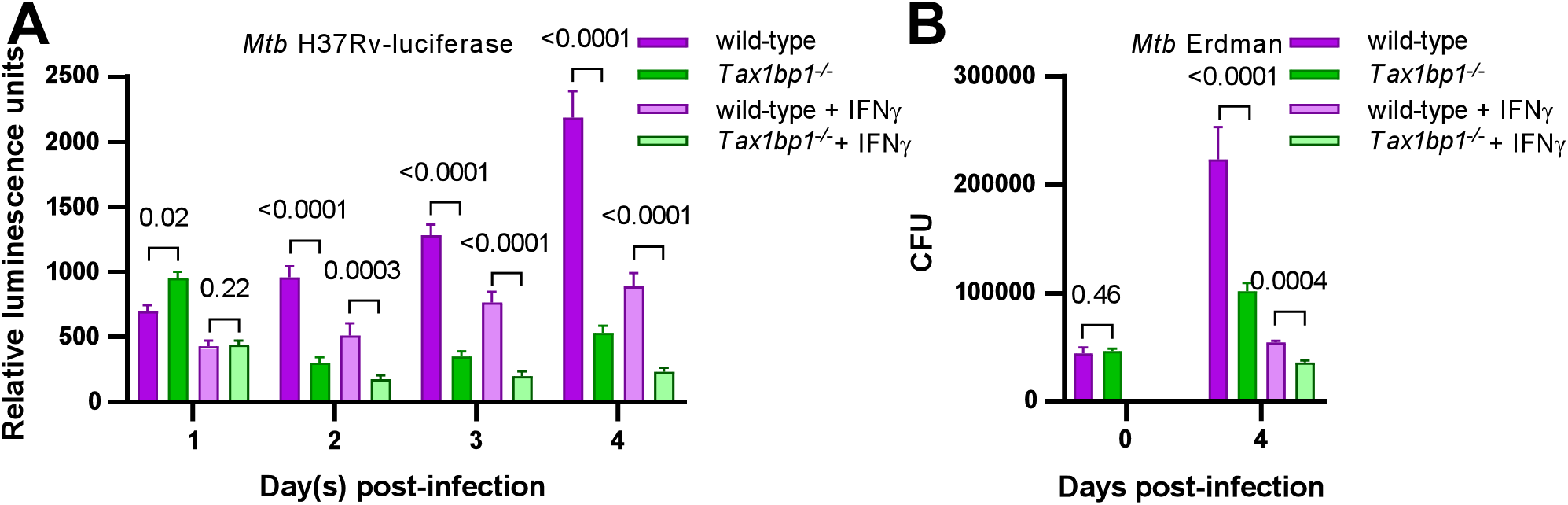
Tax1bp1 enhances *Mtb* growth in AMs infected *ex vivo*. AMs were infected *ex vivo* with luciferase-expressing *Mtb* H37Rv (A) or wild-type *Mtb* Erdman (B) at a M.O.I. of 1 in the presence or absence of IFN-γ added at the time of infection. (A) Monolayer luminescence was measured daily. (B) CFU were measured immediately after infection (day 0) or 4 days post-infection. Displayed are the mean, SEM, and FDR-adjusted p-values from the t*-*test.

### Tax1bp1 promotes *Mtb* autophagosome maturation in AMs *ex vivo*

Since we previously showed that Tax1bp1 promoted *Mtb* autophagosome maturation in BMDMs, we questioned if Tax1bp1 played a similar role in mediating selective autophagy in AMs. To test this, we performed confocal fluorescence microscopy of wild-type and *Tax1bp1^−/−^*AMs infected with fluorescent *Mtb* and assessed the colocalization of *Mtb* and autophagy markers. At 24 hours post-infection, Tax1bp1 slightly decreased *Mtb* colocalization with ubiquitin by 16% (Figure 5A-C), whereas Tax1bp1 increased *Mtb* colocalization with LC3 by 26% (Figure 5A-C). The decrease of ubiquitylated *Mtb* autophagosomes and the increase in LC3-*Mtb* colocalization indicate that Tax1bp1 promotes *Mtb* autophagosome formation and maturation in AMs to a small degree. This is the same pattern of autophagy marker staining as we previously observed in wild-type and *Tax1bp1^−/−^*BMDMs (43). Since Tax1bp1’s impact on *Mtb* growth was cell type-specific but its effect on autophagy targeting of *Mtb* was not, we reasoned that Tax1bp1 enhanced *Mtb* growth in AMs by a different mechanism.

**Figure 5.**
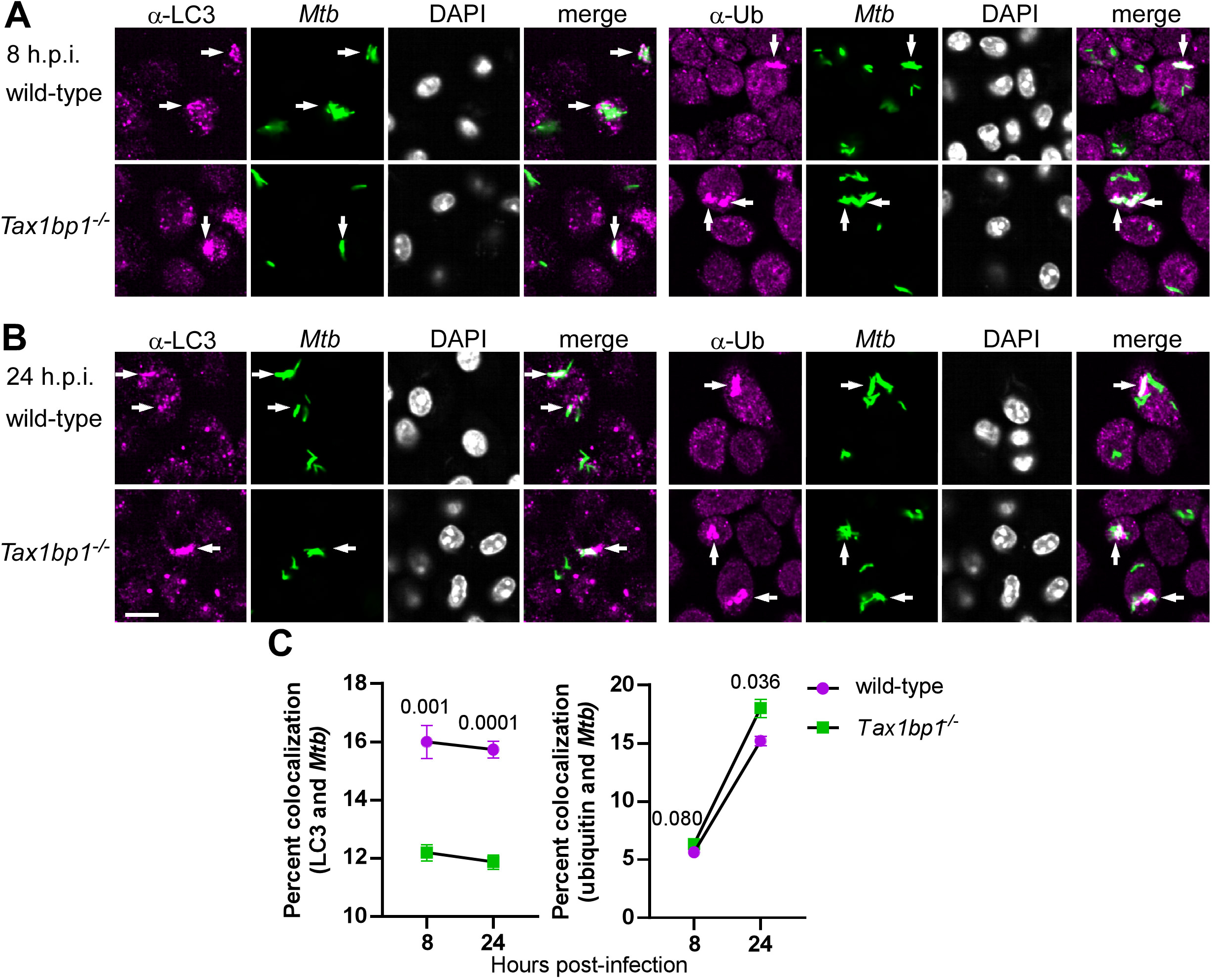
Tax1bp1 targets *Mtb* to autophagy in AMs. AMs were infected *ex vivo* with ZsGreen-expressing *Mtb* Erdman at a M.O.I of 2. At 8– and 24-hours post-infection, monolayers were fixed and stained with primary antibodies for autophagy markers, secondary Alexa-Fluor 647 antibodies, and DAPI. Immunofluorescence microscopy was performed at 63X magnification in 69 x/y positions and 4 z planes each in quadruplicate wells. Immunofluorescence microscopy images at 8-(A) and 24-(B) hours post-infection are displayed. Arrows denote *Mtb* that colocalized with LC3 or ubiquitin. The white bar denotes 10 µm. (C) Quantification of *Mtb* and autophagy marker colocalization is displayed. Mean percent colocalization in each well, SEM, and p-values from the t-test are depicted.

### Host and pathogen expression analysis of *Mtb-*infected *Tax1bp1^−/−^* AMs *ex vivo*

To broadly query host effector responses regulated by Tax1bp1 and simultaneously determine if Tax1bp1 triggers upregulation of *Mtb* genes required for intracellular *Mtb* replication, we performed host and pathogen dual transcriptional profiling of *Mtb-*infected wild-type and *Tax1bp1^−/−^* AMs. Differential gene expression analysis of *Mtb* transcripts showed no significant changes in gene expression using an adjusted p-value cutoff of 0.05. However, using a less stringent p-value threshold for statistical significance of an unadjusted p-value of 0.05, similar to other reports of *Mtb* transcriptional profiling (54), Tax1bp1 upregulated two *Mtb* genes, *mmpL4* and *mbtE* (Figure 6-figure supplement 1A). The upregulated *Mtb* genes included several required for intracellular *Mtb* replication, including *mmpL4* (55) and genes with a p-value >0.05, including the extracellular repeat protein TB18.5 (56) and transketolase *tkt* (57) (Figure 6-figure supplement 1A). These results are consistent with our observation that Tax1bp1 enhances increased intracellular *Mtb* growth during AM infection (Figure 4).

**Figure 6.**
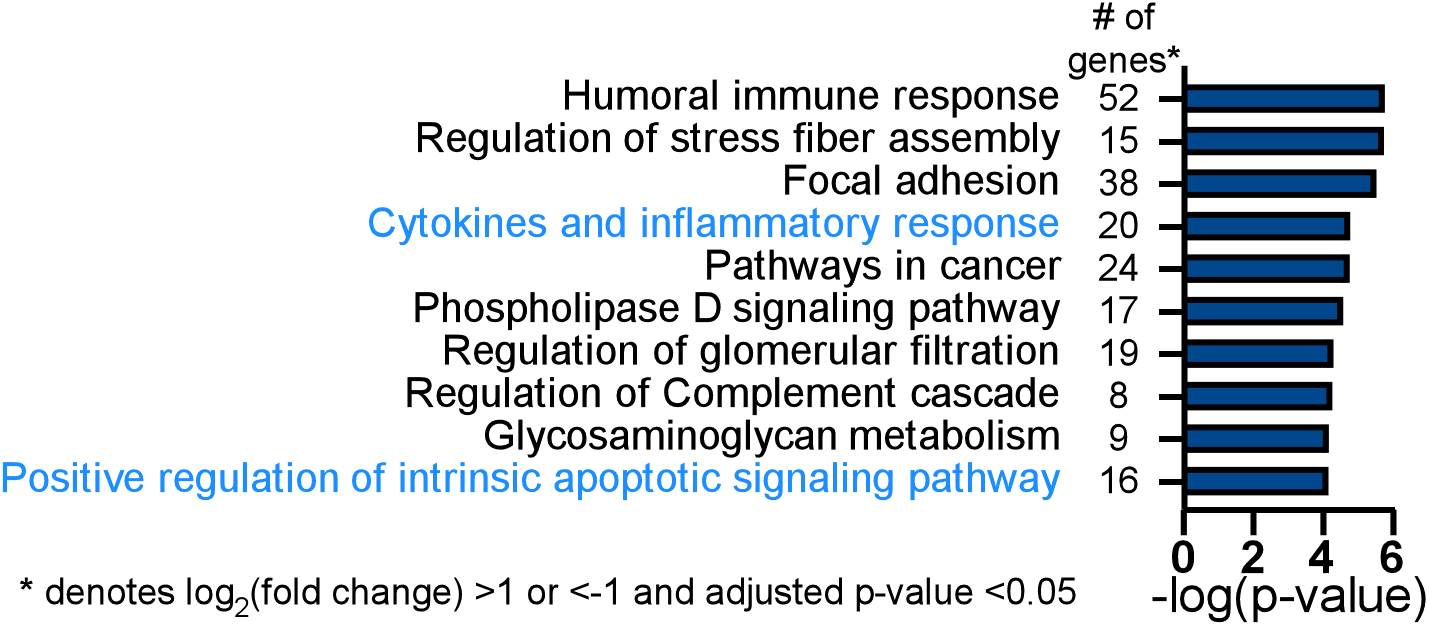
Tax1bp1 contributes to differential expression of inflammatory response and apoptotic signaling pathway genes during *Mtb* infection of AMs. Wild-type and *Tax1bp1^−/−^*AMs were infected in biological triplicate with *Mtb* at a M.O.I. of 2. The RNA was harvested at 36-hours post-infection for differential pathogen and host gene expression analysis by RNAseq. Gene ontogeny enrichment analysis of statistically significant differentially expressed host genes (log_2_(fold change) >1 or <-1, adj. p-values <0.05) during *Mtb* infection of wild-type and *Tax1bp1^−/−^* AMs was performed with Metascape (93). The top ten enriched pathways and the number of genes in each functional pathway are displayed.

From the 140 differentially expressed host genes with a log_2_(fold change) of greater than 1 or less than –1 and an adjusted p-value <0.05, gene ontogeny enrichment analysis identified cytokines and inflammatory response (20 genes, –log (p*-*value) = 4.8) as a significant functional pathway controlled by Tax1bp1 in *Mtb-*infected AMs, including the genes *Cd4*, *Cxcl1*, *Pf4*, *Pdgfa*, *Kitl*, and *Cxcl3* (Figure 6; Figure 6-figure supplement 1B). Gene ontogeny enrichment analysis also identified positive regulation of the intrinsic apoptotic signaling pathway (16 genes, –log (p*-*value) = 4.1), including the genes encoding for Bok, an apoptotic regulator, and prostaglandin-endoperoxide synthase 2 (*Ptgs2*, also known as COX-2; Figure 6-figure supplement 1B). The differentially expressed gene of the greatest magnitude in *Mtb-* infected *Tax1bp1^−/−^* AMs compared to wild-type AMs was *Sox7* (log_2_(fold change) = 5.4, adjusted p-value = 2.94 x 10^-12^). Sox7 is a transcription factor that induces apoptosis through the MAP kinase ERK-BIM (BCL2-interacting mediator of cell death) pathway (58). These results from gene expression analysis indicate that Tax1bp1 regulates the expression of inflammatory signaling and host cell death genes, both of which can contribute to the control of *Mtb* growth. Therefore, we further tested the hypothesis that Tax1bp1 impacts these two host responses during *Mtb* infection of AMs.

### Tax1bp1 promotes inflammatory cytokines signaling during AM infection *ex vivo*

We next aimed to investigate whether the effects of Tax1bp1 on cytokines and inflammatory responses during AM infection would align with our previous observation that Tax1bp1 promoted inflammatory cytokine production during infection *in vivo* (Figure 1 C, D, and Figure 1-figure supplement 1B). Tax1bp1 promoted gene expression of IL-1β (Fig. 6-figure supplement 1B) and synthesis of several inflammatory cytokines, including IL-1β, but not IFN-β as measured by ELISA (Figure 7A). As previously noted, Tax1bp1 also led to an increase in the expression of *Ptgs2* (prostaglandin-endoperoxide synthase 2; Fig. 6-figure supplement 1B), which is involved in the production of inflammatory prostaglandins.

**Figure 7.**
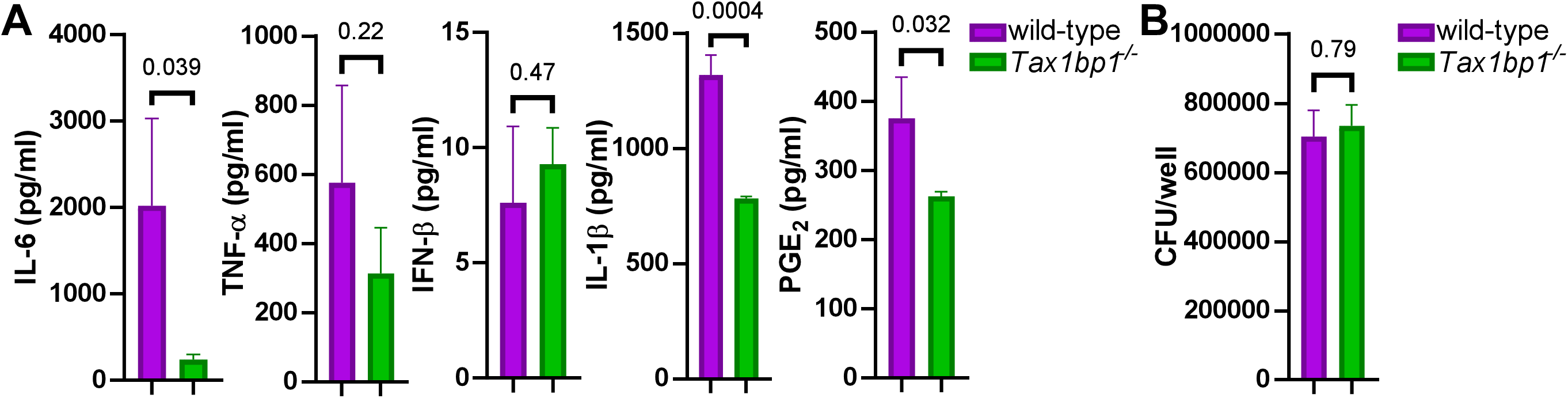
Tax1bp1 enhances IL-6, IL-1β, and PGE_2_ secretion during AM infection. AMs were seeded at 100,000 cells/well and infected in triplicate wells with *Mtb* at a M.O.I. of 5. At 24 hours post-infection, (A) the supernatants were collected for cytokine measurement by ELISA, and (B) AM monolayers were lysed and plated for *Mtb* CFU. Mean, SEM, and p-values from the t*-*test are displayed.

Notably, prostaglandin E_2_ (PGE_2_) is an inflammatory eicosanoid that promotes *Mtb* growth by blocking efferocytosis (59,60). In addition to enhancing cytokines regulated by NF-κB, Tax1bp1 also promoted the production of PGE_2_ during *Mtb* infection of AMs (Figure 7A). To our knowledge, PGE_2_ was not previously known to be regulated by Tax1bp1. Importantly, these cytokine and eicosanoid levels were measured early after infection (24 hours) when the *Mtb* CFU were the same in wild-type and *Tax1bp1^−/−^* AMs (Figure 7B), before any differences in *Mtb* growth were observed at later time points. Therefore, these findings suggest that the increased cytokine and PGE_2_ production mediated by Tax1bp1 happens independently of bacterial burden. In summary, Tax1bp1 promotes the production of proinflammatory cytokines and eicosanoids during *Mtb* infection of AMs. These findings are consistent with our cytokine analysis during *in vivo* infection (Figures 1 and 2) and in contrast to our previous report in BMDMs in which Tax1bp1 did not impact inflammatory cytokine production during *Mtb* infection (43).

### Tax1bp1 enhances necrotic-like cell death and delays apoptosis of *Mtb-*infected AMs

In addition to regulating cytokine and inflammatory responses, our gene expression analysis suggested that Tax1bp1 regulates apoptotic gene expression during AM infection. Tax1bp1 was previously shown to regulate apoptosis following cytokine stimulation (61) and viral infection (42). Furthermore, apoptosis is a mode of cell death that leads to restriction of *Mtb* growth through efferocytosis, in which *Mtb-*infected apoptotic cells are phagocytosed by neighboring macrophages (59,62). This is in contrast to necrosis, which is a form of uncontrolled cell death that is highly immunostimulatory and enhances *Mtb* growth (46,63–66). To test if Tax1bp1 impacts cell death during *Mtb* infection of AMs and BMDMs, we analyzed cells during *Mtb* infection by live cell fluorescence microscopy using CellEvent caspase 3/7 to detect apoptotic cells and propidium iodide (PI) for necrotic/late apoptotic cells. In the absence of IFN-γ stimulation, Tax1bp1 leads to necrotic-like cell death and a delay in apoptosis (Figure 8A, B; Figure 8-figure supplement 1). In IFN-γ stimulated cells, Tax1bp1 also delays apoptosis (Figure 8C, D; Figure 8-figure supplement 2). In contrast to AMs, Tax1bp1 did not impact the amount of apoptotic or necrotic-like cell death in infected BMDMs in the absence of IFN-γ (Figure 8-figure supplement 2A, C). Only when stimulated with IFN-γ, Tax1bp1 delayed apoptotic cell death during *Mtb* infection of BMDMs on day four post-infection (Figure 7-figure supplement 2B, D). Together, these results show that Tax1bp1 leads to necrotic-like cell death of AMs, but not BMDMs, and causes a delay in apoptosis during *Mtb* infection. These results also highlight an important difference in cell death modality between *Mtb-*infected wild-type AMs and BMDMs. AMs begin to undergo necrotic-like cell death early after infection, followed by apoptosis later, whereas *Mtb-*infected BMDMs also die from apoptosis later but do not undergo early necrosis.

**Figure 8.**
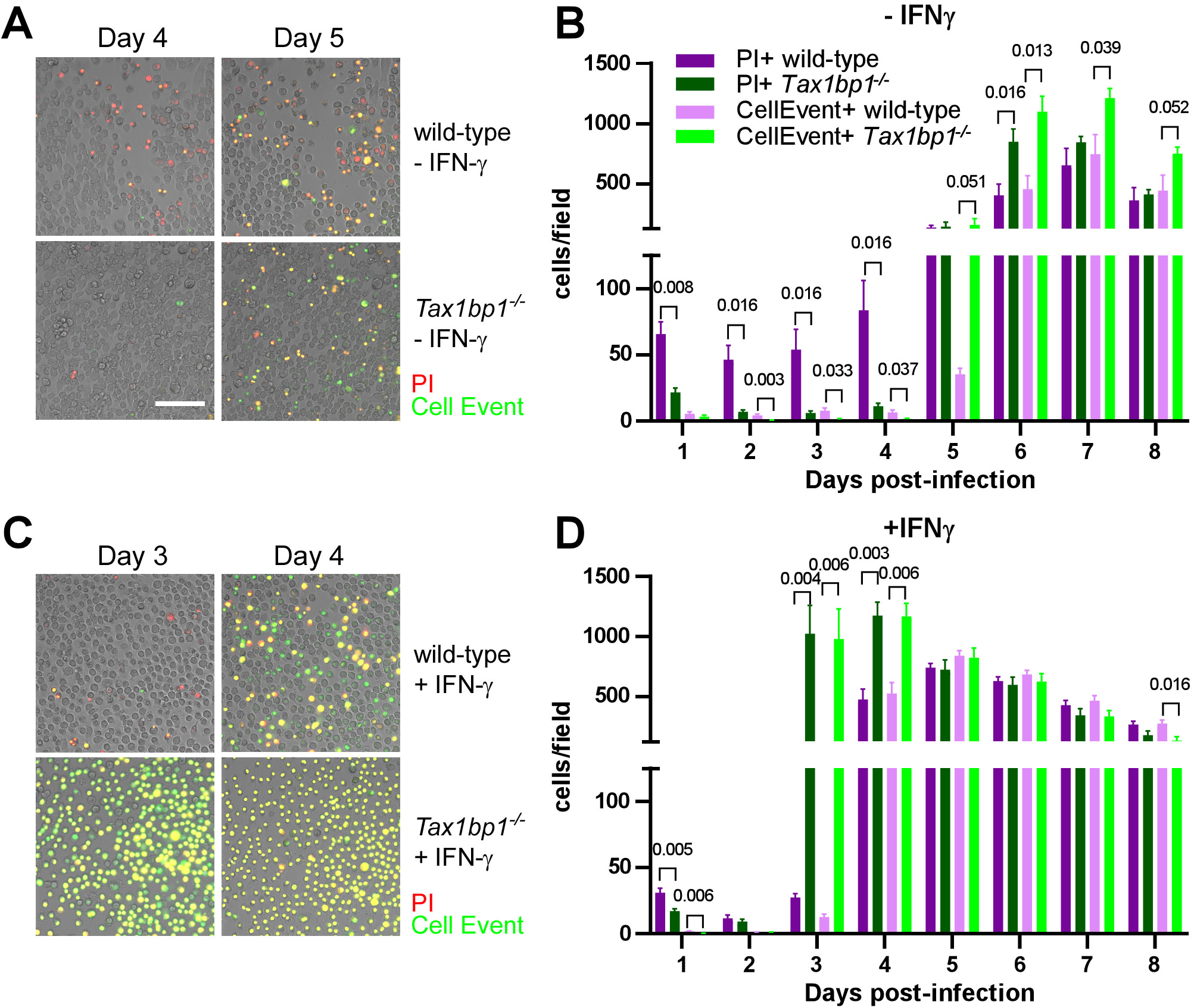
Tax1bp1 promotes necrotic-like cell death and delays apoptosis in *Mtb-*infected AMs. AMs were infected with *Mtb* at a M.O.I. of 1 in the presence of PI (propidium iodide) and CellEvent without (A, B) or with (C, D) IFN-γ added to the media. Fluorescence images were obtained at 20X magnification in two positions per well in three replicate wells. (A, C) Representative fluorescence and brightfield microscopy images were merged, cropped, and scaled. (B, D) The number of fluorescent cells in each field was quantified in the green (CellEvent) and red fluorescence (PI) channels. Mean, SEM, and statistically significant FDR-adjusted p-values from t-test comparisons are displayed. For clarity, only statistically significant p-values (p ≤ 0.05) are shown. The white bar is 100 μm.

### Expression of phosphosite-deficient Tax1bp1 restricts *Mtb* growth in AMs

We next investigated the function of Tax1bp1 phosphorylation because Tax1bp1 phosphorylation controls inflammatory responses (40). In data obtained from murine embryonic fibroblasts and *in vitro* kinase assays, Tax1bp1 is phosphorylated at 13 amino acids by the non-canonical IκB kinase IKKi (or IKK epsilon) (41), eight amino acids by the non-canonical IκB kinase TBK1 (67), and two amino acids by the canonical IKK kinase IKKα (40,41). Phosphorylation of Tax1bp1 by IKKα terminates inflammatory signaling triggered by cytokine or LPS stimulation (40). We initially discovered that Tax1bp1 is significantly phosphorylated during *Mtb* infection compared to mock infection at the IKKα substrate serine-693 in a global phosphoproteomic analysis of BMDMs (43). Having found that Tax1bp1 enhances inflammatory signaling during *Mtb* infection of AMs (Figure 7A), we hypothesized that phosphorylation of Tax1bp1 has a critical functional role during *Mtb* infection.

To test if Tax1bp1 phosphorylation impacts *Mtb* growth, we took advantage of the expanded AM (exAM) model in which primary murine AMs replicate in the presence of granulocyte-macrophage colony-stimulating factor (GM-CSF) (68,69). We initially transduced primary murine *Tax1bp1^−/−^* exAMs with lentivirus for overexpression of Flag-tagged wild-type, a double phosphomutant Tax1bp1 alleles incapable of phosphorylation by IKKα (alanine substitution; Flag-Tax1bp1^S619A, S693A^), or another that mimics phosphorylation (glutamic acid substitution; Tax1bp1^S619E, S693E^). However, we could not recover enough transduced *Tax1bp1^−/−^* exAMs for subsequent assays (data not shown). Therefore, we overexpressed the Flag-Tax1bp1 alleles in wild-type exAMs, as confirmed by immunoblot of cellular lysates with anti-Flag antibodies (Figure 9A). We then infected the transduced exAMs with *Mtb* and measured intracellular growth of *Mtb* by CFU analysis four days post-infection. Compared to wild-type exAMs transduced with empty vector, overexpression of Flag-Tax1bp1 in wild-type exAMs led to a non-significant increase in *Mtb* growth (Figure 9B). However, compared to Flag-Tax1bp1, overexpression of Flag-Tax1bp1^S619A, S693A^ but not Flag-Tax1bp1^S619E, S693E^ restricted *Mtb* growth in a statistically significant manner (Figure 9B). We acknowledge several limitations of this approach. First, Tax1bp1 phosphomutants may interact with wild-type Tax1bp1 still present in these cells. Second, the exAM model does not include all the metabolic cues found *in vivo* (*e.g.,* TGF-β) (70). In summary, these results reveal that expression of phosphosite-deficient Tax1bp1 inhibits *Mtb* growth in exAMs, similar to our finding that *Tax1bp1^−/−^* AMs also restrict *Mtb* growth compared to wild-type AMs.

**Figure 9.**
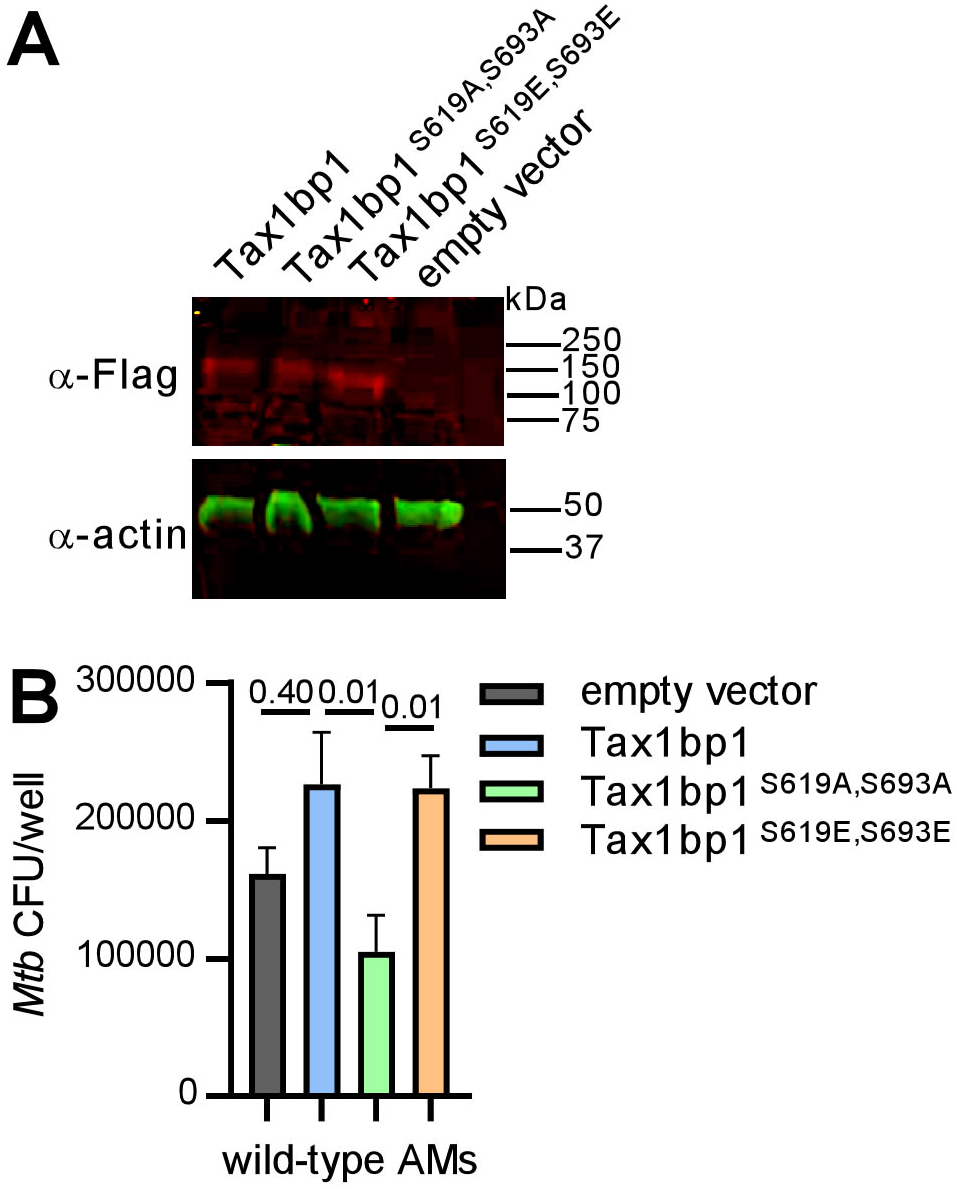
Overexpression of phosphosite-deficient Tax1bp1 restricts *Mtb* growth in AMs. (A) Cell lysates from wild-type AMs transduced with lentivirus for overexpression of Flag-tagged wild-type or phosphomutant Tax1bp1 were separated by SDS-PAGE. Immunoblot was performed with primary antibodies for the Flag epitope or actin and secondary antibodies conjugated to IRdye 680RD or IRdye 800CW, respectively. Fluorescence images in the 680 (red) and 800 (green) channels are displayed. The mobility of the molecular weight marker is displayed. (B) Transduced AMs were infected with *Mtb* (M.O.I. 0.5) in 5 technical replicate wells, and CFU enumerated at 4 days post-infection. Mean, SEM and adjusted p*-*values from Tukey’s multiple comparison test (ordinary one-way ANOVA) are displayed.

## Discussion

The autophagy receptor Tax1bp1 plays a role in multiple stages of intracellular pathogen infections, including autophagy and regulation of cytokine responses. More specifically, Tax1bp1 was implicated in the termination of inflammatory NF-κB signaling during Sendai virus and VSV infection (39) as well as the restriction of *Mtb* growth in BMDMs (43). Here we show that Tax1bp1, unexpectedly, also plays a role in promoting inflammation *in vivo* and in *Mtb* growth in AMs, neutrophils, and MNC2 (Figure 3, Figure 3-figure supplement 2). While Tax1bp1 did not impact *Listeria* growth in macrophage *ex vivo*, Tax1bp1 supports *Listeria* and *Mtb* infection in mice (Figure 2C, D, F, and Figure 2-figure supplement 1; Figure 1B, E, and Figure 1-figure supplement 1A). In addition to *Mtb* and *Listeria,* differing results between *ex vivo* and *in vivo* pathogen replication were also reported during RSV infection (37). Viral replication is restricted during *Tax1bp1^−/−^* murine infection but only slightly changed in cultured A549 *Tax1bp1* knockdown cells (37). Collectively, the differences in pathogen replication in mice and cultured cells suggest that the cell type and the tissue environment *in vivo* can play a critical role in the function of Tax1bp1. Indeed, transplanted bone marrow precursors and terminally differentiated macrophages can change their chromatin landscape in various tissue environments (71). This has significant consequences in the lungs, where the microenvironment impacts macrophage activation and function (72). The discovery that Tax1bp1 promotes *Mtb* infection in AMs, MNC2, and neutrophils, implies that Tax1bp1’s function supports *Mtb* replication in several innate immune cell types, including those from different embryonic origins. In contrast, Tax1bp1 restricts *Mtb* replication in BMDMs and, likely, MNC1. One major difference between BMDMs and AMs is that M-CSF (macrophage colony stimulating factor (73,74)) is used as a stimulating factor for the differentiation of the former, whereas GM-CSF (75–77) is thought to be more crucial for the differentiation and maintenance of the latter (78,79). Since these stimulating factors induce phenotypic changes in macrophages (80) and GM-CSF can be a bactericidal effector against *Mtb* (81), testing whether the stimulating factor present during immune cell differentiation enables Tax1bp1 to promote or restrict *Mtb* growth may shed light on an underlying mechanism that drives Tax1bp1’s cell type-specific function.

In addition to inducing inflammatory signaling, we discovered that Tax1bp1 controls the mode of host cell death by initially promoting necrotic-like cell death in the first four days of *Mtb* infection in AMs but not BMDMs, while delaying apoptosis in the later stages of infection. Since *Mtb* growth is enhanced in necrotic macrophages (46,82) and apoptosis leads to restriction of *Mtb* growth via efferocytosis (59), these results indicate that Tax1bp1’s impact on the mode of host cell death is a mechanism by which Tax1bp1 enhances *Mtb* growth in AMs. Indeed, Tax1bp1 was previously shown to mediate ferroptosis, a programmed type of necrosis, in response to copper stress-induced reactive oxygen species (83). Interestingly, ferroptosis enhances *Mtb* dissemination (84). In addition to promoting ferroptosis, Tax1bp1 is known to impact apoptotic signaling by restraining apoptosis during VSV and Sendai virus infection (42). During viral infection, termination of RIG-I mediated mitochondrial antiviral signaling proteins (MAVS) signaling blocked apoptosis and type I IFN signaling (42). Because we did not observe altered levels of type I IFN during AM infection with *Mtb*, we hypothesize Tax1bp1 signals through a different pathway during *Mtb* infection in these cells. Tax1bp1 also blocks apoptosis by acting as an adaptor for TNFAIP3 (also known as A20) to bind and inactivate its substrates RIPK1 (receptor (TNFRSF)-interacting serine-threonine kinase 1) in the TNFR signaling pathway (85). Further experiments are needed to determine the downstream signaling pathway by which Tax1bp1 blocks apoptosis and whether it promotes a programmed form of necrosis during *Mtb* infection.

Tax1bp1 is an autophagy adaptor, and autophagy plays a role in cell-autonomous immunity to microbial pathogens. Autophagy also affects immune cell development and inflammatory responses (86–90). The *Tax1bp1*-deficient knockout mice used in this study are deficient in *Tax1bp1* in all cells. While our results indicate that the effect of Tax1bp1 is mediated during the innate immune responses, other cells may likely require Tax1bp1 for their function and that might impact *Mtb* infection. For example, Tax1bp1 is important for the metabolic transition of activated T cells (91). Tax1bp1 also terminates ERK signaling in B cells to mediate B cell differentiation and antigen-specific antibody production (92). Therefore, Tax1bp1 may be needed for the normal function of other immune cells that undergo proliferation and activation in addition to AMs.

In conclusion, Tax1bp1 plays a unique role in controlling host cell death and promoting inflammatory responses during *Mtb* and *Listeria* infection in contrast to its function in terminating NF-κB signaling during viral infection. We discovered that Tax1bp1 is a host factor contributing to differences in *Mtb* growth in AMs compared to BMDMs (8). While multiple autophagy receptors, including Tax1bp1 and p62, target *Mtb* for selective autophagy (22,43), this work reveals that different autophagy receptors may play fundamentally distinct roles in *Mtb* pathogenesis even depending on the host cell type. In contrast to Tax1bp1, the primary autophagy receptor, p62, is not involved in survival from *Mtb* infection (48). These findings would suggest that one could alter the inflammatory responses and augment protective host responses to pathogens by blocking Tax1bp1 function or inhibiting Tax1bp1 phosphorylation. A better understanding of the mechanism by which Tax1bp1 regulates host cell death, autophagy, and inflammation during infection may enable the development of Tax1bp1 as a target for anti-bacterial therapies.

## Supplemental Figure Legends

**Figure 1-figure supplement 1.**
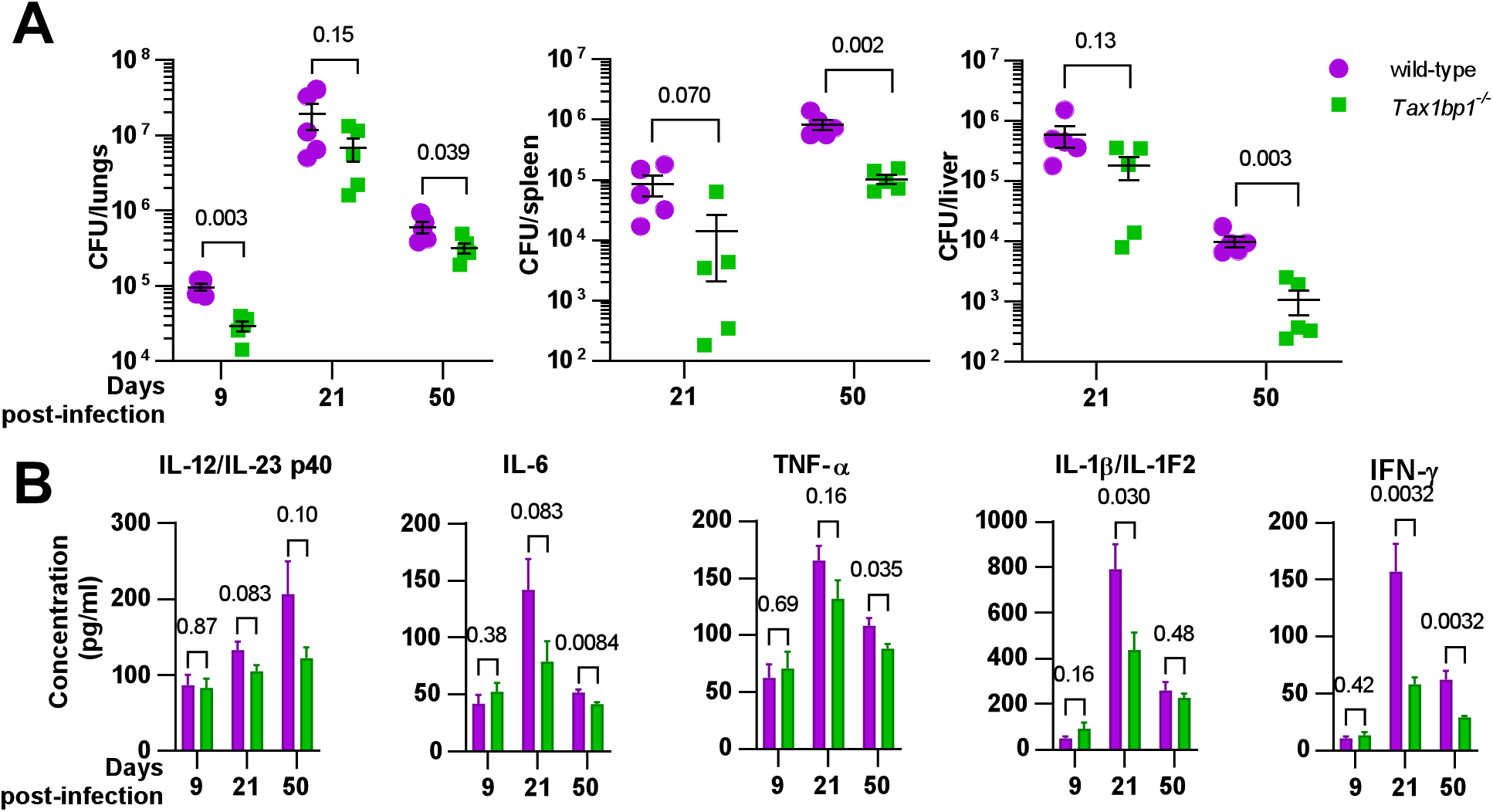
Tax1bp1 contributes to *M. tuberculosis* virulence and inflammatory cytokine responses. (A) In an independent experiment, male and female mice infected by the aerosol route with *M. tuberculosis* were euthanized at 1-, 9-, 21-, and 50 days post-infection. Lung homogenates were enumerated for CFU. Results are the mean ± SEM from five mice. The mean experimental inoculum was 104 CFU as determined by CFU enumeration at 1-day post-infection. (B) Cytokine levels from infected lung homogenates at 9-, 21-, and 50 days post-infection were measured by ELISA. Results are the mean ± SEM from five samples. The p-values from t-test comparisons are shown.

**Figure 1-figure supplement 2.**
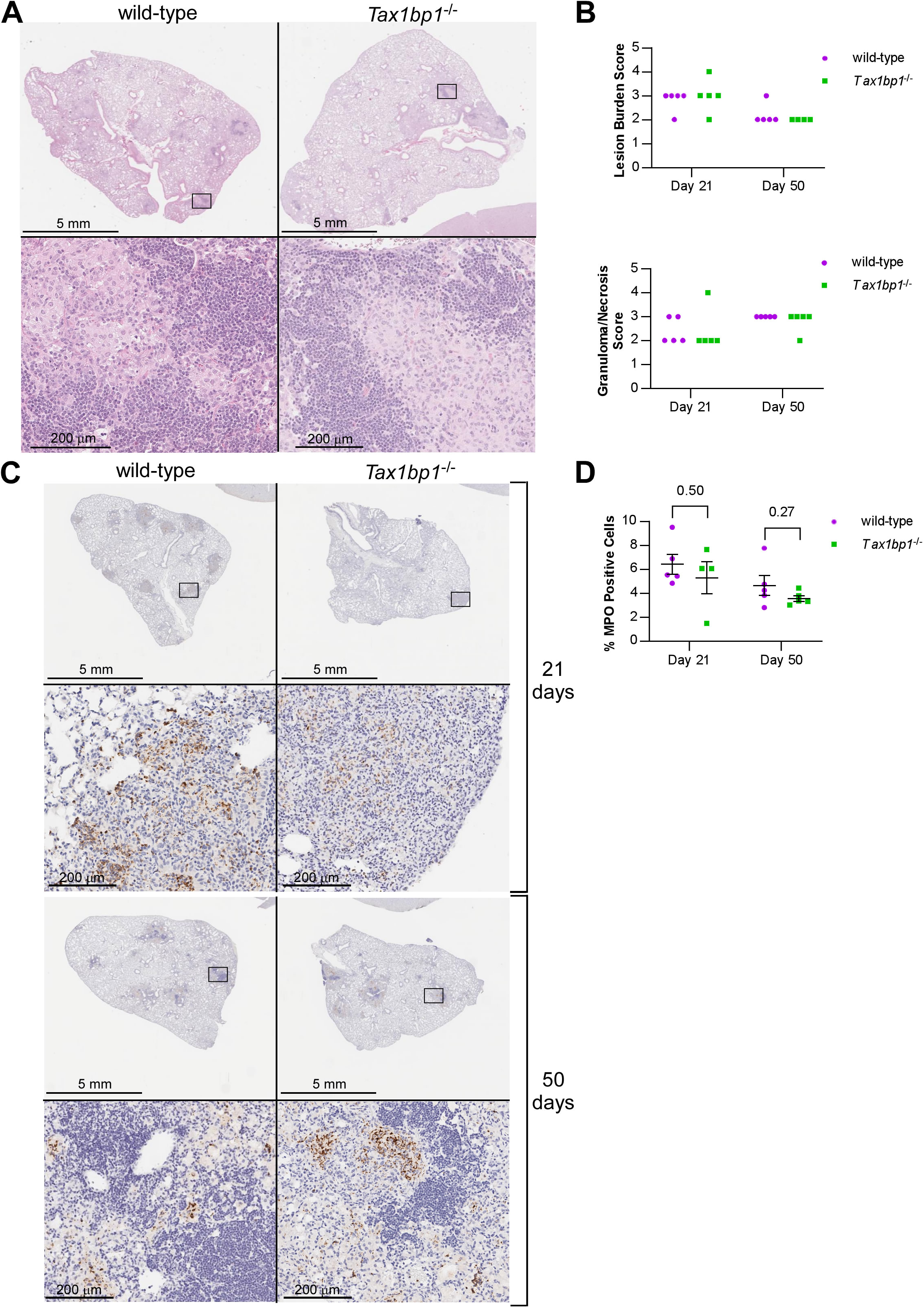
Lung pathology and neutrophil recruitment were similar during *M. tuberculosis* aerosol infection of wild-type and *Tax1bp1^−/−^* mice. (A) Paraffin-embedded thin sections of lung samples from infected wild-type and *Tax1bp1^−/−^* mice were stained with hematoxylin and eosin (H&E). (B) Pathology was analyzed in H&E-stained images from five infected wild-type and five *Tax1bp1^−/−^*mice at 21– and 50 days post-infection. (C) Paraffin-embedded thin sections of the lung from infected wild-type and *Tax1bp1^−/−^* mice were stained with antibodies against myeloperoxidase. Antibody staining was detected with 3,3’-diaminobenzidine. (D) Quantitative analysis of the percentage of cells that stained positive for myeloperoxidase is shown. Results are the mean ± SEM from five mice. Brackets indicate p-values from t-test comparisons.

**Figure 1-figure supplement 3.**
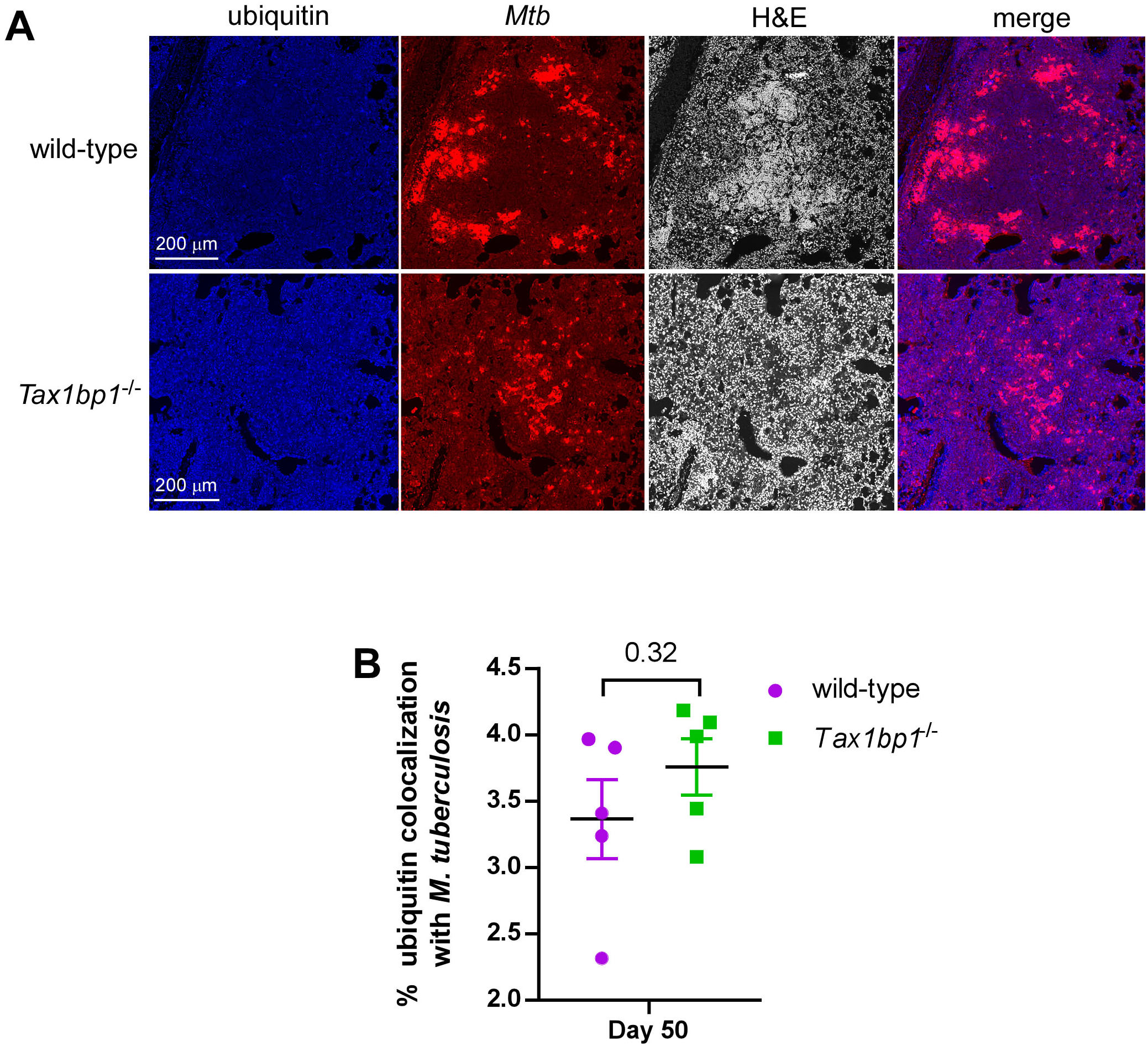
Ubiquitin colocalizes with *M. tuberculosis* in the lungs during murine aerosol infection of wild-type and *Tax1bp1^−/−^* mice. (A) Serial thin sections of paraffin-embedded lung specimens were stained with antibodies against ubiquitin, *M. tuberculosis*, or hematoxylin and eosin. Antibodies were detected with 3,3’-diaminobenzidine. (B) Quantitative analysis of ubiquitin staining pixel overlap with *M. tuberculosis* in overlayed images. Results are mean ± SEM from five samples. The p*-*value from the t-test comparison is shown.

**Figure 2-figure supplement 1.**
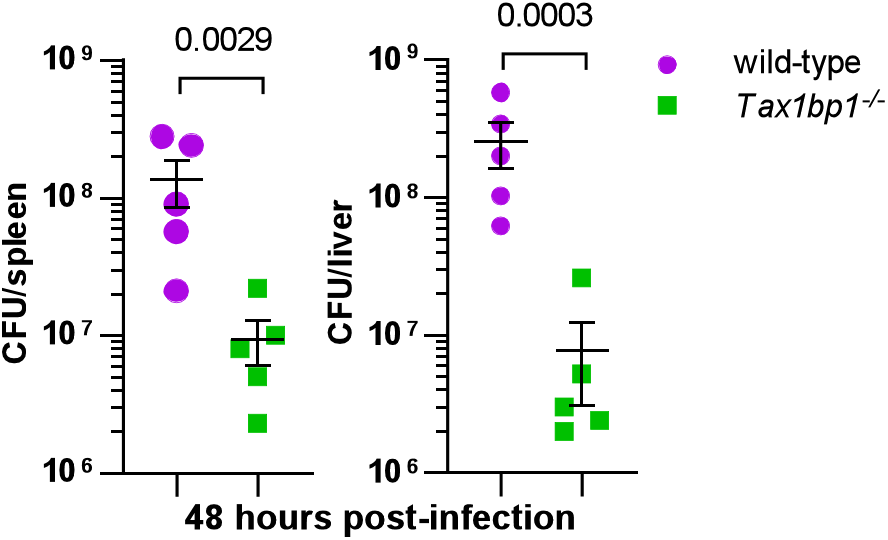
Tax1bp1 enhances *L. monocytogenes* growth during murine infection. In an independent experiment, mice were infected with *L. monocytogenes* by the intravenous route, and CFU enumerated from spleen and liver homogenates at 48 hours post-infection. Results are the mean ± SEM from five mice. Brackets indicate p-values from t-test comparisons. CFU data were logarithmically transformed prior to statistical analysis.

**Figure 2-figure supplement 2.**
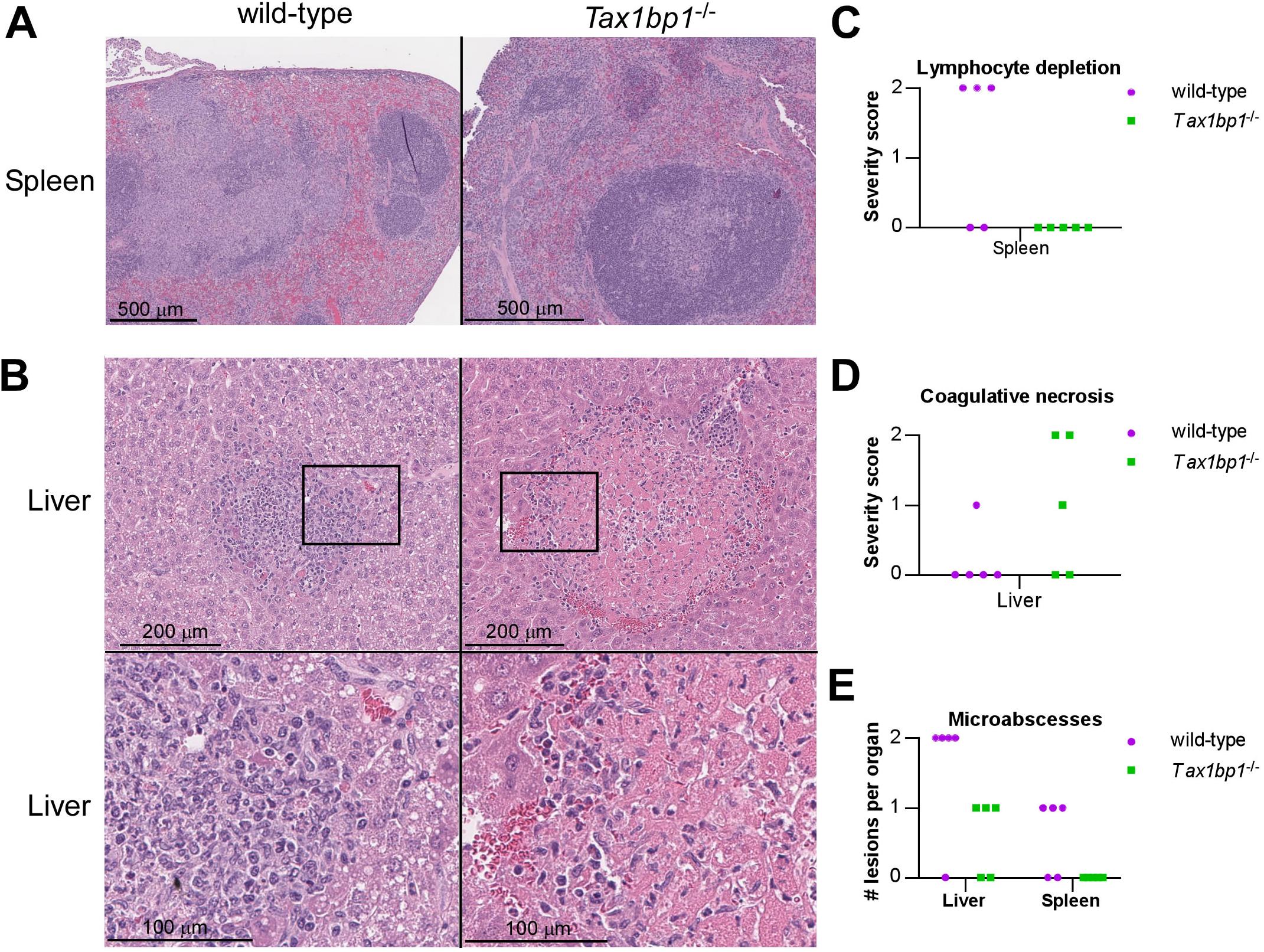
Tax1bp1 promotes the formation of microabscesses and lymphocyte depletion during *L. monocytogenes* infection. (A, B) Serial thin sections of paraffin-embedded spleen and liver specimens from mice infected by the intraperitoneal route collected at 72 hours post-infection were stained with hematoxylin and eosin. (C-E) Pathology was analyzed in H&E-stained images from five infected wild-type and 5 *Tax1bp1^−/−^* mice at 72 hours post-infection.

**Figure 3-figure supplement 1.**
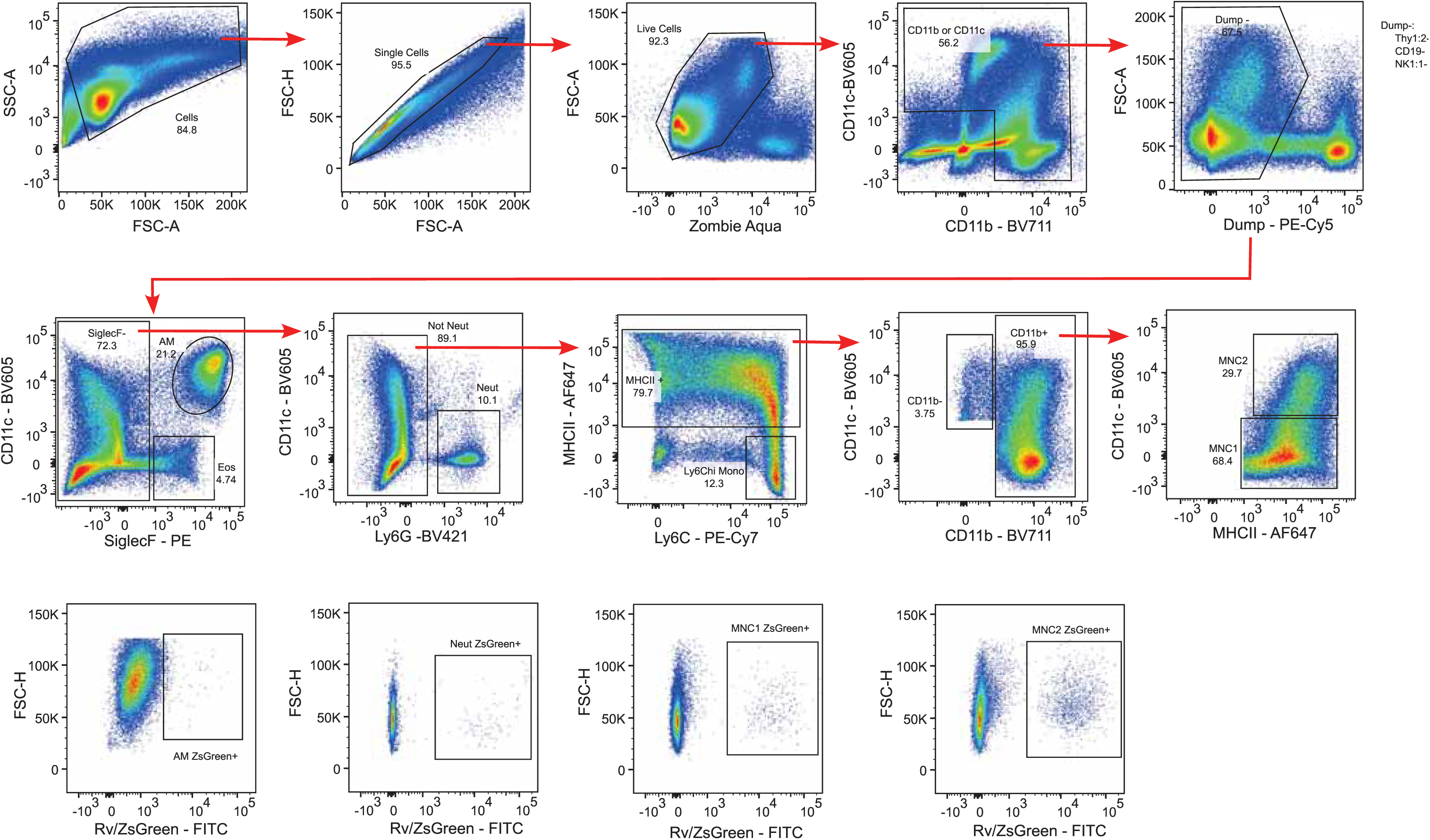
Gating strategy used to identify myeloid subsets. A representative flow panel is shown depicting the gating strategy for identification and sorting of myeloid subsets. B, T, and NK cells were gated out. AMs (CD11b^lo^CD11c^hi^SiglecF^hi^), MNC1 (SiglecF^-^CD11b^+^CD11c^lo^MHCII^+^), MNC2 (SiglecF^-^CD11b^+^CD11c^hi^MHCII^hi^), and neutrophils (Neut; SiglecF^-^Ly6G^hi^CD11b^hi^) were sorted. Sorted cells were plated for *Mtb* CFU enumeration. The gating strategy used to identify ZsGreen-positive cells is shown in the bottom row.

**Figure 3-figure supplement 2.**
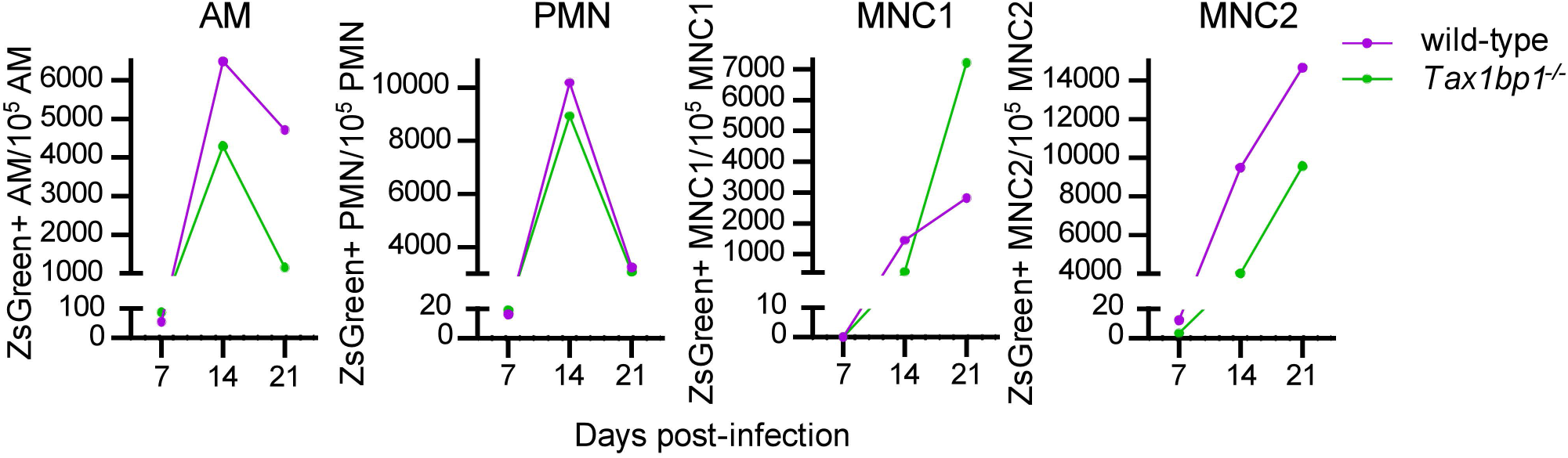
Tax1bp1 promotes *Mtb* growth in AMs, PMNs, and MNC2 following low-dose aerosol infection. In an independent experiment, mice were infected with aerosolized *Mtb* expressing ZsGreen. Five wild-type and five *Tax1bp1^−/−^* mice were euthanized at 1-, 7-, 14-, and 21 days post-infection. CFU were measured from lung homogenates at 1 day post-infection, which revealed the mean infectious dose of *Mtb* was 224 CFU/mouse. At 7-, 14-, and 21 days post-infection, lung cells were pooled and stained for AMs, PMNs, and recruited monocyte 1 and 2 subsets (MNC1, 2). The number of ZsGreen-positive counts from innate immune cells was quantified by analytical flow cytometry. Data were normalized to the number of cells analyzed.

**Figure 6-figure supplement 1.**
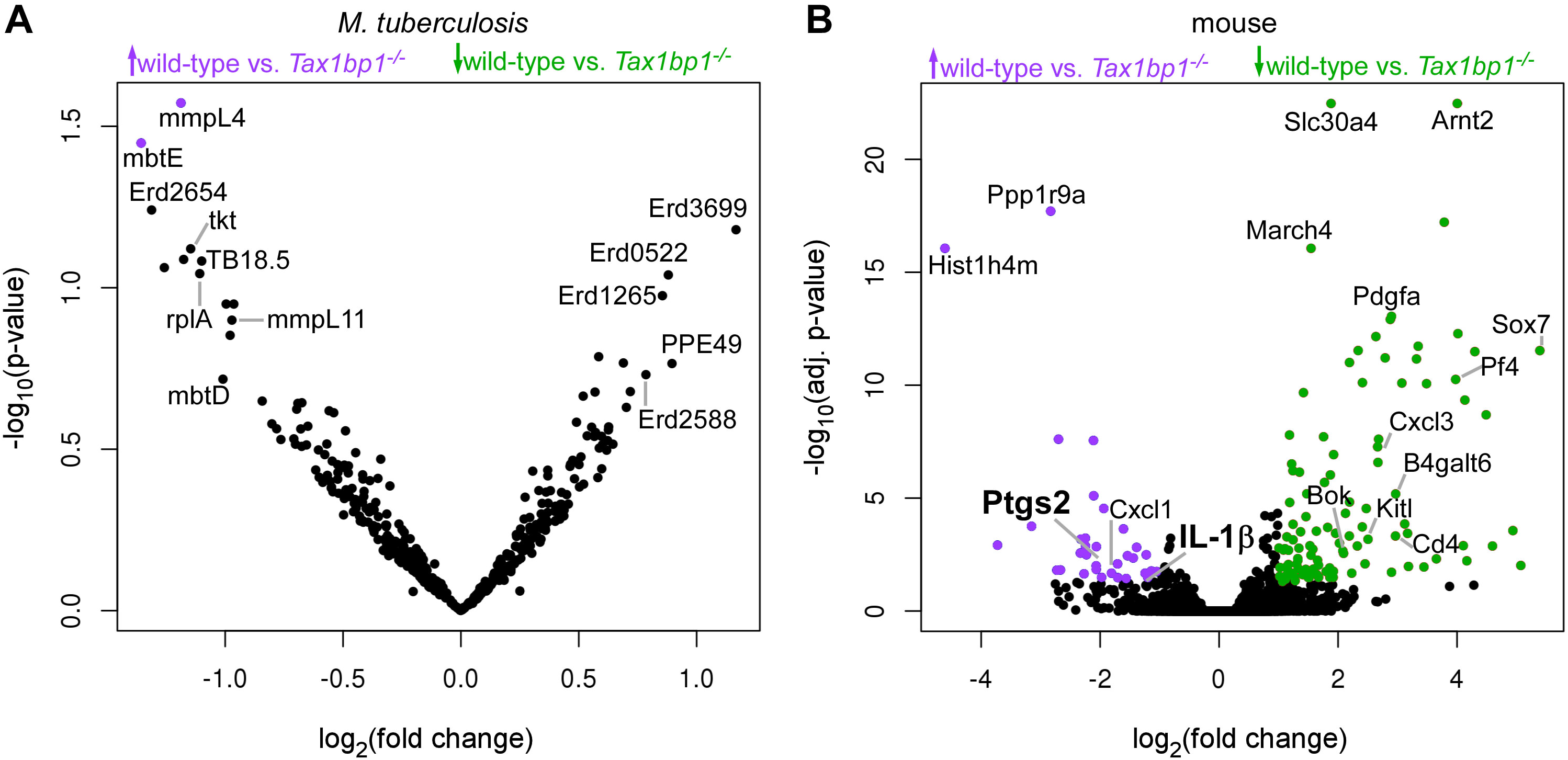
Pathogen and host differential gene expression analysis volcano plots. Volcano plots display the differentially regulated genes from (A) *Mtb* and (B) the host during wild-type and *Tax1bp1^−/−^*AM infection with *Mtb*. The volcano plots display the log_2_fold change of normalized mean hit counts in wild-type vs. *Tax1bp1^−/−^*samples and –log_10_(adj. p-value for host genes or unadjusted p*-*value for *Mtb* genes). Colors denote genes that were upregulated (purple) or downregulated (green) in wild-type compared to *Tax1bp1^−/−^* samples.

**Figure 8-figure supplement 1.**
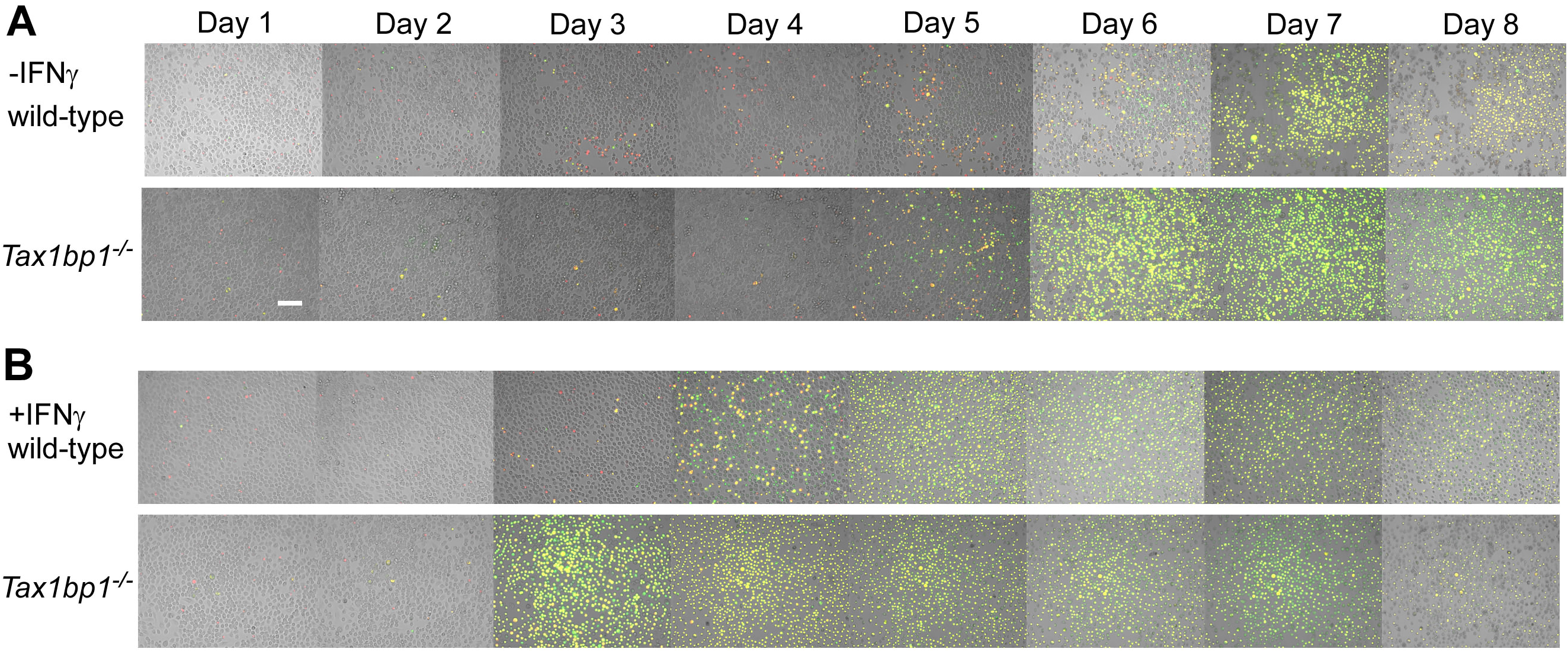
Tax1bp1 enhances necrotic-like cell death and delays apoptosis during *Mtb* infection of AMs. As described in the Figure 8 legend, AMs were infected with *Mtb* at a M.O.I. of 1 in the presence of PI (propidium iodide) and CellEvent without (A) or with (B) IFN-γ added to the media. Fluorescence images were obtained at 20X magnification in two positions per well in three replicate wells. Representative fluorescence and brightfield microscopy images are displayed at days 1-8 post-infection. The white bar is 100 µm.

**Figure 8-figure supplement 2.**
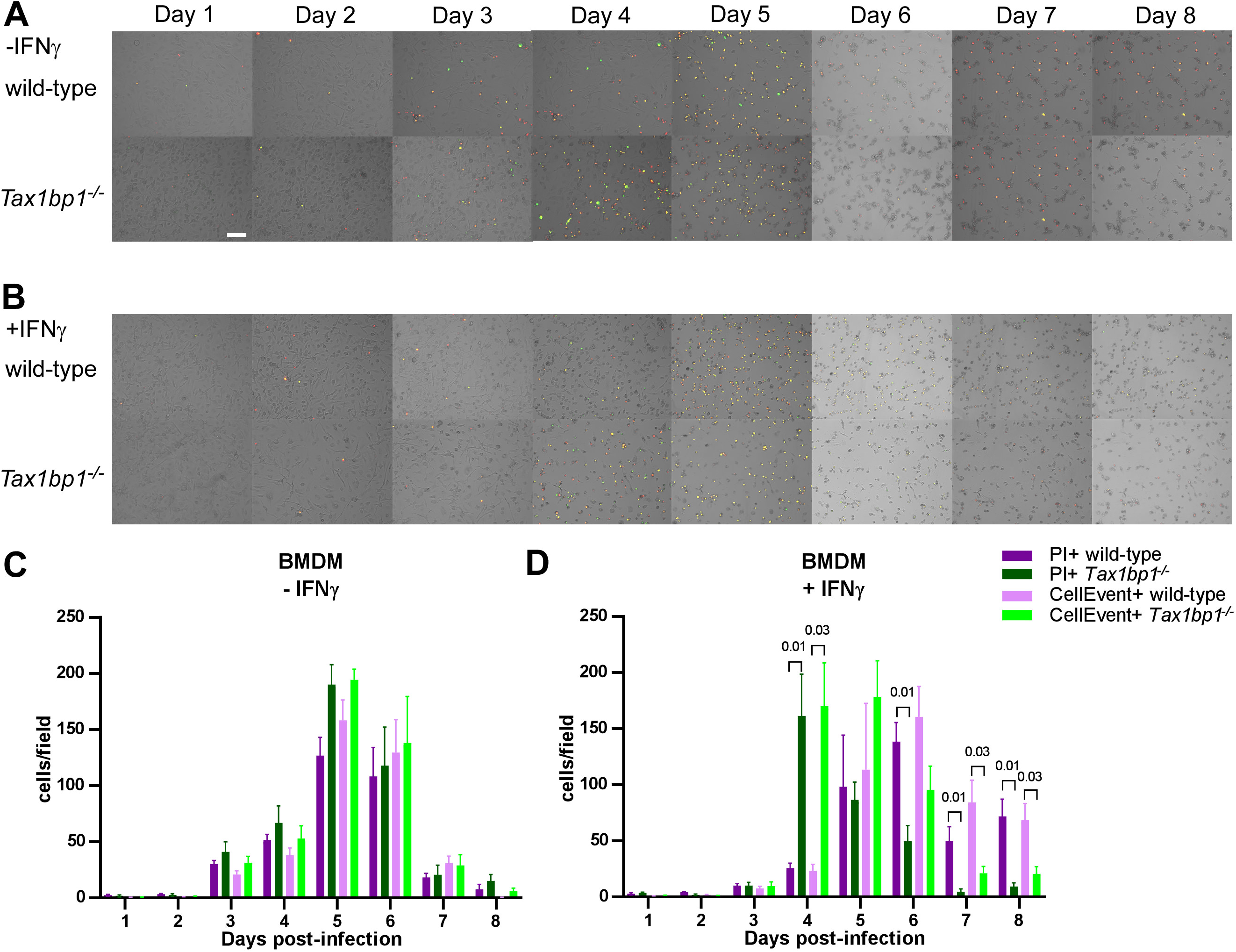
Tax1bp1 does not promote necrotic-like cell death during *Mtb* infection of BMDMs. BMDMs were infected with *Mtb* at a M.O.I. of 1 in the presence of PI (propidium iodide) and CellEvent without (A, C) or with (B, D) IFN-γ added to the media. Fluorescence images were obtained at 20X magnification in two positions per well in three replicate wells. Representative fluorescence and brightfield microscopy images are displayed at days 1-8 post-infection. The white bar denotes 100 µm. (C, D) The number of fluorescent cells in each field was quantified in the green (CellEvent) and red fluorescence (PI) channels. Mean, SEM, and statistically significant FDR-adjusted p-values comparisons are displayed. For clarity, only statistically significant p-values (p ≤ 0.05) are displayed.

**Figure 8-figure supplement 1.**
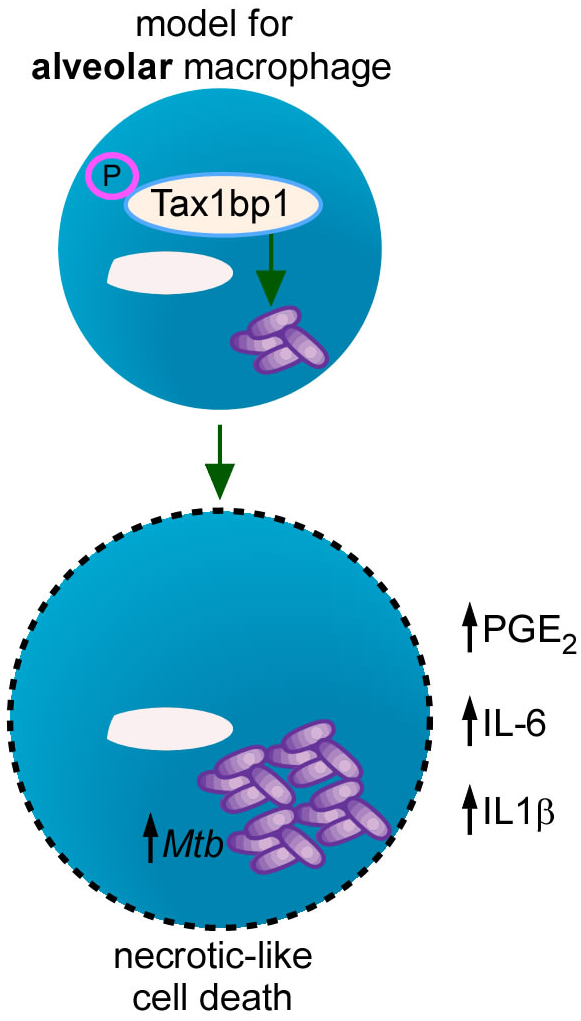
Model describing Tax1bp1’s function during *Mtb* infection of AMs. Tax1bp1 enhances *Mtb* growth, inflammatory cytokine synthesis, PGE_2_ production, and necrotic-like host cell death in AMs. Tax1bp1-deficiency, or expression of phosphosite-deficient Tax1bp1, decreases *Mtb* growth in AMs.

## Methods

### Ethics statement

Animal infections were performed in accordance with the animal use protocol (AUP-2015-11-8096, AN192778-01) approved by the Animal Care and Use Committee at the University of California, Berkeley, and the Institutional Animal Care and Use Program at the University of California, San Francisco, in adherence with the federal regulations provided by the National Research Council and National Institutes of Health.

### *M. tuberculosis* mouse infections at UC Berkeley

*Tax1bp1^−/−^* mice were provided by Dr. Hidekatsu Iha, Oita University, Japan. Low-dose aerosol infection (100 CFU) of age– and sex-matched wild-type or Tax1bp1-deficient mice (male and female, age 8-12 weeks) was performed with *M. tuberculosis* Erdman strain using the Glass-Col Inhalation Exposure System. One day after infection, infected mice were euthanized, the lungs were homogenized, and CFU were enumerated on 7H10 agar plates supplemented with 10% OADC and 0.5% glycerol to determine the inoculum. On days 9, 11, 21, and 50 days after infection, the lung was divided into portions. The superior lobe of the right lung was fixed in 10% buffered formalin for histologic analysis. The remainder of the right lung and left lung were combined and homogenized in 1 ml of PBS containing 0.05% Tween80 in a Bullet Blender Tissue Homogenizer (Next Advance). The spleen and liver were homogenized in 400 µl or 2 ml of PBS containing 0.05% Tween80, respectively. For measurement of CFU, organ homogenates were serially diluted and plated on 7H10 agar plates supplemented with 10% OADC and 0.5% glycerol. For measurement of cytokines from the lung homogenate, protein extraction was performed by combining 700 µl of homogenate with 700 µl of Tissue Protein Extraction Reagent (T-PER; Thermo Scientific) containing cOmplete, mini, EDTA-free protease inhibitor cocktail (Roche 11836170001) at 2X concentration. Samples were incubated for 20 min at 4°C, vortexed, and centrifuged at 12,000 x g for 10 min. The supernatants were filter sterilized with a 0.22 μm filter and stored at –80°C until further analysis. For survival experiments, mice were sacrificed after 15% loss of maximum body weight.

### Cytokine measurements

Cytokines from lung lysates were measured using DuoSet ELISA kits (R&D systems) following the manufacturer’s protocol. Interferon levels were measured from the *Mtb-*infected lung lysates using L929 ISRE-luciferase reporter cells as previously described (94). Luciferase reporter cells were seeded in a 96-well plate for 24 hours. Lung lysates were incubated with the reporter cells for 8 hours, and luciferase activity was measured with the Luciferase Assay Report Assay (Promega) using the manufacturer’s protocol.

### Histology sample processing and quantitative analysis

Formalin-fixed specimens were washed three times in PBS and stored in 70% ethanol. Histologic processing was performed by Histowiz. Serial ultrathin sections were stained for hematoxylin & eosin (H&E), ubiquitin (anti-ubiquitylated antibody, AB1690, EMD-Millipore, 1:100 dilution), tuberculosis (Abcam ab214721, 1:1000 dilution), or myeloperoxidase (Abcam ab9535, 1:50 dilution). Primary antibodies were detected with 3,3’-diaminobenzidine (DAB) staining. The ubiquitin and tuberculosis IHC images were aligned and combined using image registration scripts in QuPath. The MPO staining analysis was analyzed using Indica Labs Halo image analysis software. Cells were segmented using the Multiplex IHC algorithm v3.1.4 in Halo and MPO-positive cells were determined by thresholding the DAB channel. Positive cells were further sub-divided into Low intensity, Medium Intensity and High intensity bins to allow for subsequent calculation of a H-score based on percentage of cells positive for Low, Medium and High using the following formula: H-Score = (1 x % positive cells low) + (2 x % positive cells medium) + (3 x % positive cells high). Cells in all the intensity bins were considered positive for myeloperoxidase staining.

Images of tuberculosis lesions were exported from QuPath into ImageJ. A minimum threshold of two standard deviations above the mean signal was applied to filter positive pixels for ubiquitin and tuberculosis immunohistochemistry images. Colocalization was calculated from pixel overlap in images of the tuberculosis and ubiquitin immunohistochemistry. A veterinary pathologist analyzed the H&E histopathology images.

### *M. tuberculosis* mouse infections at UC San Francisco

*Tax1bp1^−/−^* mice were imported from UC Berkeley and rederived by the UC San Francisco Rederivation Core to eliminate the potential for any interinstitutional murine pathogen transmission. Low-dose aerosol infections (100-200 CFU) of age– and sex-matched wild-type and *Tax1bp1^−/−^* mice were performed with a Glass-col inhalation chamber. At the indicated time points, mice were euthanized with CO_2_, and their lungs were minced with scissors and digested in 3 ml of RPMI-1640 with 5% heat-inactivated FBS containing 1 mg/ml collagenase D (Sigma) and 30 µg/ml DNAseI (Sigma) for 30 min at 37°C. Cells were processed with a gentleMACS dissociator (Miltenyi Biotec, lung program 2) and filtered through a 70 μm strainer. The samples were rinsed with 1 ml of FACS buffer (PBS with 3% heat-inactivated FBS, 2 mM EDTA). Residual tissue on the cell strainer was further processed using a syringe plunger and rinsed with 1 ml of FACS buffer. The cell suspension was then centrifuged at 650 ˣ g for 3 minutes at 4 °C, and the supernatant was discarded. The cell pellet was resuspended in 3 ml of ACK lysis buffer (Gibco) to lyse the RBCs, and lysis was quenched with 3 ml of FACS buffer solution. After centrifuging the cell suspension at 650 ˣ g for 3 minutes at 4 °C, the supernatant was removed, and the cells were resuspended in 1 ml of FACS buffer. Each cell suspension was pooled (from 5 mice) and passed through a 50-µm strainer.

Single-cell lung suspensions were stained with the Zombie Aqua Fixable Viability kit (1:200 dilution, BioLegend, #423101) and treated with CD16/CD32 Fc block (1:100 dilution, BD 553142) in PBS (1 ml) for 15 min. The samples were centrifuged at 650 ˣ g for 3 minutes at 4 °C, and the supernatants were removed. Cells were stained with 2 ml of antibody mixture diluted in Brilliant Stain Buffer (Table 1; Invitrogen #00-4409-42) for 30 min at 4°C. Antibodies diluted in Brilliant Stain Buffer (BD, #566349) were added to the cells. Antibody staining was performed for 30 minutes at 4°C. Subsequently, the cell suspensions were centrifuged at 650 g for 3 minutes at 4°C, washed with 1 ml of FACS buffer solution, resuspended in 3 ml of FACS buffer, and passed through a 50-μm strainer. Cell subsets were sorted using a BD Aria Fusion Sorter through a 100 µm nozzle using the 4-way purity mode.

**Table 1.** Key Resources.

| Reagent type (species) or resource | Designation | Source or reference | Identifiers | Additional information |
| --- | --- | --- | --- | --- |
| Biological sample (*M. musculus*) | Primary bone marrow-derived macrophages, AMs, and peritoneal exudate cells, wild-type C57BL/6J | Jackson Laboratory | Stock # 000664 |  |
| Biological sample (*M. musculus*) | Primary bone marrow-derived macrophages, AMs, and peritoneal exudate cells, *Tax1bp1*<sup>-/-</sup> | (36) | *Tax1bp1*<sup>-/-</sup> |  |
| Antibody | Anti-ubiquitylated antibody | EMD-Millipore | AB1690 | 1:100 dilution |
| Antibody | Anti-*Mycobacterium tuberculosis* antibody | Abcam | ab214721 | 1:1000 dilution |
| Antibody | Anti-Myeloperoxidase antibody | Abcam | ab9535 | 1:50 dilution |
| Antibody | Anti-LC3 (2G6) | NanoTools | 0260-100/LC3-2G6 | 1:200 |
| Antibody | Anti-ubiquitinated proteins antibody (FK2) | Sigma-Aldrich | 04-263 | 1:400 |
| Antibody | PE Rat Anti-Mouse Siglec-F | BD Biosciences | BDB552126 | 1:200 |
| Antibody | Anti-mouse CD16/CD32 Fc block | BD Biosciences | BD553142 | 1:100 |
| Antibody | PE/Cyanine5 anti-mouse CD90.2 (Thy1.2) | BioLegend | 105314 | 1:300 |
| Antibody | PE/Cyanine5 anti-mouse CD19 | BioLegend | 115510 | 1:300 |
| Antibody | PE/Cyanine5 anti-mouse NK-1.1 | BioLegend | 108716 | 1:200 |
| Antibody | PE/Cyanine7 anti-mouse Ly-6C | BioLegend | 128018 | 1:300 |
| Antibody | Brilliant Violet 421 anti-mouse Ly-6G | BioLegend | 127628 | 1:200 |
| Antibody | Brilliant Violet 605 anti-mouse CD11c | BioLegend | 117334 | 1:200 |
| Antibody | Brilliant Violet 711 anti-mouse/human CD11b | BioLegend | 101242 | 1:200 |
| Antibody | Alexa Fluor 647 anti-mouse MHCII | BioLegend | 107618 | 1:300 |
| Commercial assay | Luciferase assay system | Promega | E1500 |  |
| Commercial kit | Mouse IL-6 DuoSet ELISA | R and D | DY406 |  |
| Commercial kit | Mouse TNF-α DuoSet ELISA | R and D | DY410 | |
| Commercial kit | Mouse IL-12/IL-23 p40 allele-specific DuoSet ELISA | R and D | DY499 |  |
| Commercial kit | Cytometric Bead Array Mouse Inflammation Kit | BD Biosciences | 552364 |  |
| Commercial kit | Mouse IFN-β ELISA Kit, high sensitivity | PBL Assay Science | 42410-1 | |
| Commercial kit | Prostaglandin E<sub>2</sub> Express ELISA Kit | Cayman Chemicals | 500141 |  |
| Commercial kit | Q5 Site Directed Mutagenesis kit | New England Biolabs | E0554S |  |
| Commercial kit | NEBuilder HiFi DNA Assembly Cloning Kit | New England Biolabs | E5520S |  |
| Commercial kit | LR Clonase II Enzyme Mix | Invitrogen | 11791020 |  |
| Strain | Lenti-X 293T | TakaraBio | 632180 |  |
| Strain | *Mtb*: Erdman | ATCC | 35801 |  |
| Strain | *Listeria monocytogenes* 10403S | (99) |  |  |
| Strain | *Mtb*: H37Rv (pMV306hsp-LuxG13) | Plasmid from Addgene | 26161 |  |
| Strain | *Mtb*: Erdman (pMV261::ZsGreen) | (10) |  |  |

Bacteria were quantified from sorted cells by serial dilution in PBS containing 0.05% Tween80 and plated on 7H10 agar plates supplemented with 10% Middlebrook OADC, 0.5% glycerol, and PANTA antibiotic mixture at a 1:500 dilution to reduce contamination risk from non-mycobacteria during organ dissection. BD PANTA antibiotic mixture (BD, B4345114) containing polymyxin B, amphotericin B, nalidixic acid, trimethoprim, and azlocillin was prepared by dissolving the contents of 1 lyophilized vial in 3 ml of OADC.

### Bone marrow-derived macrophage infection

Bone marrow-derived macrophage infections with *Listeria monocytogenes* 10403S were performed as previously described (49–51).

### Peritoneal cell exudate infection

Approximately 7 ml of ice-cold PBS was injected into the peritoneum of euthanized mice. Peritoneal exudate cells were treated with ACK lysis buffer, resuspended in tissue culture cell media (RPMI supplemented with 1 mM L-glutamine and 10% fetal bovine serum), and seeded into 24-well plates with glass coverslips at a density of 1.5 ˣ 10^6^ cells/ml for 24-hours prior to infection. Prior to infection, non-adherent cells were removed by replacement of the tissue culture media.

*Listeria monocytogenes* 10403S was inoculated from a single colony into BHI media. Following overnight incubation at 30°C, bacteria were washed in PBS and resuspended to an OD of 1.5 in PBS. Bacteria were diluted 1:1000 in tissue culture cell media for infection of peritoneal exudate cells. 30 minutes post-infection, cells were rinsed twice with PBS, and fresh media was replaced. At 1 hour post-infection, gentamicin sulfate was added at a final concentration of 50 µg/ml to kill extracellular bacteria. At 2– and 8 hours post-infection, coverslips were placed in 5 ml of water and vortexed. Serial dilutions were plated on LB agar supplemented with streptomycin (200 µg/ml). CFU were enumerated after 24 hours of incubation at 37°C. Infections were performed with 3 coverslips for each experimental condition.

### L. monocytogenes mouse infections

Age and sex-matched male and female mice were infected with *L. monocytogenes*. A 2 ml overnight culture of *L. monocytogenes* grown in brain heart infusion media at 30°C slanted. *L. monocytogenes* was subcultured in 5.5 ml of BHI media and incubated with shaking at 37°C until reaching an optical density between 0.4-0.8. The bacteria were washed and diluted in PBS to achieve an inoculum of approximately 5 ˣ 10^5^ CFU/ml. 200 μl of this suspension was injected into the tail vein of the mice. For the intraperitoneal infection, the bacterial cells were prepared as described, and 200 µl of bacterial suspension was injected into the peritoneum at a dose of 3.74 ˣ 10^5^ CFU/mouse. At the indicated time points, the spleen and liver were harvested in water containing 0.1% NP-40. Organs were homogenized, and CFU were enumerated on LB agar supplemented with streptomycin (200 µg/ml).

### Alveolar macrophage (AM) isolation and culture

AMs were harvested from mice by bronchoalveolar lavage with 10 ml of PBS containing 2 mM EDTA, 0.5% fetal bovine serum (FBS) pre-warmed to 37°C as described previously (68,69). AMs were seeded at a density of 100,000 cells/well in 96-well plates. For short-term cultivation up to 4 days, AMs were cultured in RPMI-1640 medium supplemented with 10% (v/v) FBS, 2 mM GlutaMAX, 10 mM HEPES and 100 U ml–1 penicillin–streptomycin. After allowing at least 2 hours for adhesion, the media was replaced with fresh media without antibiotics. For cultivation >4 days, AMs were cultured in RPMI-1640 medium supplemented with 10% (v/v) FBS, 2 mM GlutaMAX, 1 mM sodium pyruvate, and 2% (v/v) GM-CSF supernatant produced by a B16 murine melanoma cell line.

### Macrophage infections with *Mtb*

*Mtb* H37Rv strain was transformed with pMV306hsp-LuxG13 for expression of *LuxCDABE*. Logarithmic phase cultures of *Mtb* (H37Rv-Lux or wild-type Erdman strain) were grown in 7H9 media supplemented with 10% Middlebrook OADC, 0.5% glycerol, 0.05% Tween80 in inkwell bottles at 37°C with rotation at 100 rpm. *Mtb* cell pellets were washed twice with PBS followed by centrifugation for 5 min at 1462 ˣ g and sonication to remove and disperse clumps*. Mtb* was resuspended in RPMI with 10% horse serum. Media was removed from the macrophage monolayers, the bacterial suspension was overlaid, and centrifugation was performed for 10 min at 162 ˣ g. Following infection, the media was replaced with cultivation media with 15 ng/ml IFN-γ (Peprotech) or without IFN-γ. In experiments performed with luminescent *Mtb,* luminescence measurements were obtained daily following media changes daily using a GloMax microplate reader (Promega). For CFU measurements, the monolayers were washed with PBS, lysed in PBS with 0.05% Tween80, serially diluted, and spread on 7H10 agar plates supplemented with 10% Middlebrook OADC and 0.5% glycerol. CFU were enumerated after 21 days of incubation at 37°C.

### Gene expression analysis during *Mtb* infection of AMs

AMs were infected with wild-type *Mtb* Erdman at a M.O.I. of 2. At 36-hours post-infection, the monolayers were washed with PBS, and the AMs were lysed in 200 µl of Trizol reagent. Samples were pooled from four technical replicate wells. The experiment was performed independently three times (*i.e.* three independent biological replicates). *Mtb* was centrifuged, the supernatant containing host RNA was removed, and the *Mtb* pellet was resuspended in 400 µl of fresh Trizol and 0.1 mm zirconia/silica beads. *Mtb* was mechanically disrupted with the Mini Bead-Beater Plus (Biospec Products) as previously described (95). 70% of the sample containing host RNA was pooled with the sample containing *Mtb* RNA, the samples were treated with 200 µl of chloroform, and RNA was purified with the Trizol Plus RNA purification kit (Ambion). Purified total RNA was treated with DNAseI (ThermoFisher) and dried by rotary evaporation in RNA stabilization tubes (Azenta US, Inc.; South Plainfield, NJ, USA). Sample QC, dual rRNA depletion for bacteria and mouse, library preparation, Illumina sequencing (2×150 bp; 30M reads per sample), and differential gene expression analysis were performed by Azenta Life Sciences US, Inc.

#### Sample QC

Total RNA samples were quantified using Qubit 2.0 Fluorometer (Life Technologies, Carlsbad, CA, USA) and RNA integrity was checked with 4200 TapeStation (Agilent Technologies, Palo Alto, CA, USA).

#### Library Preparation and Sequencing

ERCC RNA Spike-In Mix kit (cat. 4456740) from ThermoFisher Scientific was added to normalized total RNA prior to library preparation following manufacturer’s protocol. rRNA depletion was performed using QIAGEN FastSelect rRNA Bacteria + HMR Kit or HMR/Bacteria (Qiagen, Germantown, MD, USA), which was conducted following the manufacturer’s protocol. RNA sequencing libraries were constructed with the NEBNext Ultra II RNA Library Preparation Kit for Illumina by following the manufacturer’s recommendations. Briefly, enriched RNAs are fragmented for 15 minutes at 94 °C. First strand and second strand cDNA are subsequently synthesized. cDNA fragments are end repaired and adenylated at 3’ends, and universal adapters are ligated to cDNA fragments, followed by index addition and library enrichment with limited cycle PCR. Sequencing libraries were validated using the Agilent Tapestation 4200 (Agilent Technologies, Palo Alto, CA, USA), and quantified using Qubit 2.0 Fluorometer (ThermoFisher Scientific, Waltham, MA, USA) as well as by quantitative PCR (KAPA Biosystems, Wilmington, MA, USA).

The sequencing libraries were multiplexed and clustered onto a flowcell on the Illumina NovaSeq instrument according to manufacturer’s instructions. The samples were sequenced using a 2×150bp Paired End (PE) configuration. Image analysis and base calling were conducted by the NovaSeq Control Software (NCS). Raw sequence data (.bcl files) generated from Illumina NovaSeq was converted into fastq files and de-multiplexed using Illumina bcl2fastq 2.20 software. One mis-match was allowed for index sequence identification.

#### Data Analysis

After investigating the quality of the raw data, sequence reads were trimmed to remove possible adapter sequences and nucleotides with poor quality using Trimmomatic v.0.36. The trimmed reads were mapped to the *Mus musculus* and *Mtb* Erdman strain reference genomes available on ENSEMBL using the STAR aligner v.2.5.2b. BAM files were generated as a result of this step. Unique gene hit counts were calculated by using feature Counts from the Subread package v.1.5.2. Only unique reads that fell within exon regions were counted.

Using DESeq2, a comparison of gene expression between the groups of samples was performed. The Wald test was used to generate p-values and Log2 fold changes. Genes with adjusted p-values < 0.05 and absolute log_2_ fold changes >1 were called as differentially expressed genes for each comparison.

### Cytokine analysis of *Mtb-*infected AMs

AMs were infected with wild-type *Mtb* Erdman strain at a MOI of 10. At 24 hours post-infection, the supernatants were filtered through a 0.2 μm syringe filter and analyzed by ELISA for IFN-β (PBL Assay Bioscience), TNF-α, and IL-1β (R&D systems) as previously described, and prostaglandin E_2_ (Cayman Chemicals).

### Live cell imaging

AMs were infected with *Mtb* at a MOI of 1, and 0.1 µg ml^−1^ of propidium iodide (LifeTechnologies) and two drops per milliliter of CellEvent Caspase-3/7Green ReadyProbes reagent (Invitrogen) were added to the media at the beginning of the infection to measure necrosis/late apoptosis and apoptosis, respectively. Fluorescence and phase contrast images were obtained at 20x magnification with a Keyence BZ-X 700 microscope. Images were obtained daily in three technical replicate wells per condition and at two positions in each well. Quantification of the number of necrotic and apoptotic cells was performed with ImageJ version 1.54f as described previously (46). Images were converted to 8-bit (grayscale), binarized, and enumerated using the analyze particles module (size threshold 0.001-infinity).

### Immunofluorescence microscopy

AMs were infected with fluorescent *Mtb* at a MOI of 2. At 8– and 24-hours post-infection, monolayers were washed with PBS, fixed with 4% PFA for 20 minutes, washed with PBS, and stained with anti-LC3 or anti-ubiquitin primary antibodies and AlexaFluor-647 conjugated secondary antibodies as previously described (43). Images were obtained at 63x magnification from quadruplicate wells per condition, in 69 x/y positions, and 4 z positions (0 µm, 0.5 µm, 1 µm, and 1.5 µm) with an Opera Phenix microscope (Perkin Elmer). Colocalization analysis of LC3, ubiquitin, and *Mtb* was performed with Harmony version 4.9 (Perkin Elmer) using the following analysis parameters. The four z stack images in each x/y position were processed into a maximum projection. Nuclei were identified in the DAPI channel using Method B with a common threshold of 0.07 and an area threshold of > 20 µm^2^. Cytoplasm was identified in the AlexaFluor 647 channel using Method A with an individual threshold of 0.06. The find spot module was used to identify LC3 or ubiquitin “spots” in the AlexaFluor 647 channel using method C with a contrast setting of 0.42, uncorrected spot to region intensity of 3.8, and default radius. *Mtb* were identified in the AlexaFluor 488 channel using the find spot module method B with a detection sensitivity of 0.5 and splitting sensitivity of 0.5. To identify *Mtb* that colocalized with LC3 or ubiquitin “spots”, the select population module was used for the *Mtb* population with the select by mask method. The percent colocalization was calculated for each well from all the images obtained in each well using the evaluation module.

### Lentiviral transduction of AMs for Tax1bp1 phosphomutant expression

Tax1bp1 was amplified by PCR from murine cDNA using the primers named Tax1bp1 fwd and rev (Table 2) and inserted by ligation-independent cloning (NEBuilder Builder HiFi DNA Assembly Cloning Kit, NEB #E5520S) into pENTR1A no ccdB (w48-1; Addgene #17398) previously modified to express N-terminal 3x Flag-tagged proteins (96). Site-directed mutagenesis was performed to engineer alanine or glutamic acid substitution mutations in Tax1bp1 using the primers listed in Table 2 and the Q5 site-directed mutagenesis kit (New England Biolabs). Open reading frames (ORFs) were transferred into the pLENTI CMV Puro DEST (w118-1) vector using the Gateway LR Clonase II enzyme mix (Invitrogen #11791020). Lenti-X 293T cells (Takara) were transfected with lentiviral packaging vector psPAX2 (Addgene #12260), envelop vector pMD2.G (Addgene #12259), and Flag-tagged Tax1bp1 or empty destination vector (pLenti CMV Puro DEST (w118-1) as previously described (97). The supernatant containing lentivirus was filtered in a 0.45 µm syringe filter. AMs harvested from mice were infected with lentivirus by centrifugation at 1000 ˣ g for 30 min at 32°C. Transduced AMs were allowed to recover and expand for 7 days prior to harvesting with ESGRO Complete Accutase containing 1 mM EGTA (98). AMs were seeded at a density of 100,000 cells per well in 96-well plates and infected with *Mtb* at a M.O.I. of 0.5 the subsequent day. Media was changed daily. Four days post-infection, monolayers were washed with PBS, lysed, and plated for *Mtb* CFU.

**Table 2.**
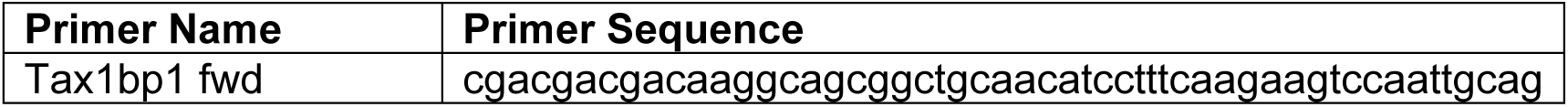
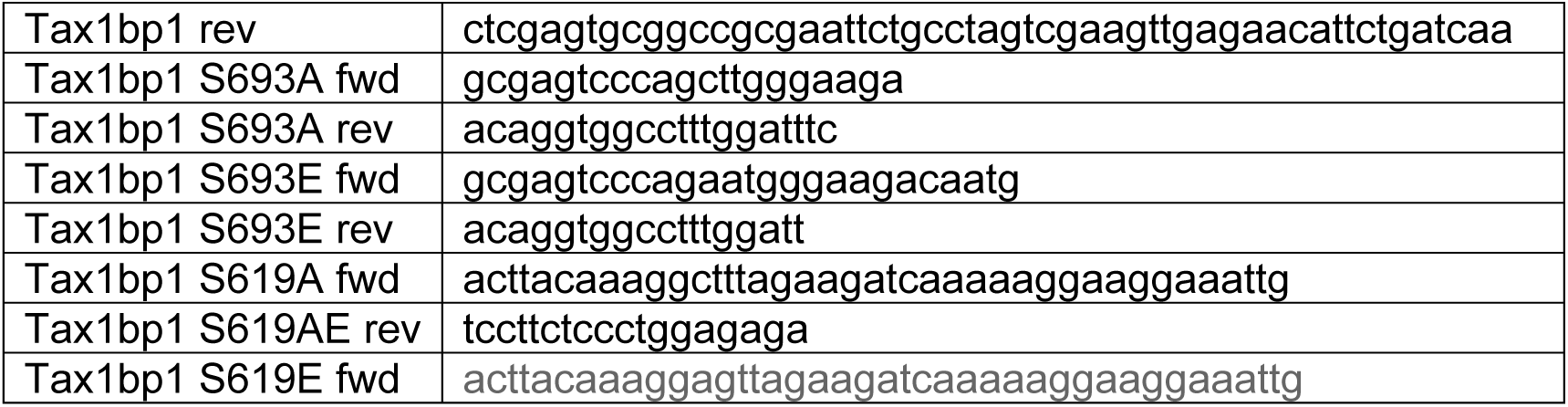
Primers used in this study.

## Data availability statement

Confocal microscopy images are available on the Dryad repository (DOI: 10.5061/dryad.44j0zpcq6). RNA sequencing files, including the differential gene expression analysis, can be accessed on GEO, accession GSE280399. Flow cytometry data files are available upon request.

## Statistical analysis

GraphPad Prism (v.10.3.1) was used for statistical analysis. Unpaired t-test comparisons were calculated assuming Gaussian distributions and the p-values were reported. In experiments with more than two experimental conditions, p-values from the t-test comparison between two groups were adjusted for the FDR (multiple comparisons) using the two-stage linear step-up procedure of Benjamini, Krieger, and Yekutieli (Figure 4, Figure 8, Figure 8-figure supplement 2). In experiments with more than two experimental conditions, the comparison between multiple groups was performed by ordinary one-way ANOVA with Tukey’s multiple comparisons test, and the adjusted p-values were reported (Figure 9).

## Acknowledgments

We acknowledge support from Professor Daniel A. Portnoy for infections with *Listeria monocytogenes.* This work was supported by NIH grants K08 AI146267 (JB), U19 AI135990 (NJK, JSC), P01 AI063302 (JSC), U19 AI106754 (JSC), DP1 AI124619 (JSC), and R01 AI120694 (JSC). JMB was also supported by the UCSF Nina Ireland Program in the Health Award, Cystic Fibrosis Foundation Harry Shwachman Award, Mentored Scientist in Tuberculosis Award (R25AI147375), and TB RAMP program (R25AI147375). R.R.-L was supported by a grant from the National Academies of Sciences, Engineering, and Medicine (Ford Foundation Fellowship) and the University of California Dissertation-Year Fellowship.

